# The RNA Structurome in the Asexual Blood Stages of Malaria Pathogen *Plasmodium falciparum*

**DOI:** 10.1101/2020.10.25.354316

**Authors:** Diana Renteria Alvarez, Alejandra Ospina, Tiffany Barwell, Bo Zheng, Abhishek Dey, Chong Li, Shrabani Basu, Xinghua Shi, Sabah Kadri, Kausik Chakrabarti

**Affiliations:** Department of Biological Sciences, University of North Carolina at Charlotte, Charlotte, North Carolina 28223; Department of Computer & Information Sciences, College of Science and Technology Temple University, Philadelphia, PA 19122; Division of Medical genetics, Children’s Hospital of Pittsburgh of UPMC, 4401 Penn Avenue Pittsburgh, PA 15224; Division of Heath and Biomedical Informatics, Northwestern University Feinberg School of Medicine and Ann & Robert H. Lurie Children’s Hospital of Chicago, Chicago, IL 60611

## Abstract

RNA as an effector of biological functions often adopts secondary and tertiary structural folds. *Plasmodium falciparum* is a deadly human pathogen responsible for the devastating disease called malaria. In this study, we measured the differential accumulation of RNA secondary structures in coding and noncoding transcripts from the asexual developmental cycle in *P. falciparum* in human red blood cells. Our comprehensive analysis, combining high-throughput nuclease mapping of RNA structures by duplex RNA-seq, immunoaffinity purification and RNA analysis, collectively measured differentially base-paired RNA regions during the parasite development. Our mapping data not only aligned to a diverse pool of RNAs with known structures but also enabled us to identify new structural RNA regions in the malaria genome. On average, ~71% of the genes with secondary structures are found to be protein coding mRNAs. Mapping pattern of these base-paired RNAs corresponded to all parts of protein-coding mRNAs, including 5’ UTR, CDS and 3’ UTR. In addition to histone family genes which are known to form secondary structures in their mRNAs, transcripts from genes which are important for transcriptional and post-transcriptional control, such as unique plant-like transcription factor family, *ApiAP2*, DNA/RNA binding protein family, *Alba*, ribosomal proteins and eukaryotic initiation factors involved in translational control and the ones important for RBC invasion and cytoadherence also show strong accumulation of duplex RNA reads in various asexual stages. Intriguingly, our study determined a positive relationship between mRNA structural contents and translation efficiency in *P. falciparum* asexual blood stages, suggesting an essential role of RNA structural changes in malaria gene expression programs.

## Introduction

RNA is innately structured. The dynamic intra- and inter-molecular interactions of RNA govern almost all steps of gene expression programs, from transcriptional activation to RNA processing and translation (Sharp 2009). Changes in RNA folding affect biological functions in a cell in a measurable fashion and are facilitated by several factors, such as temperature, ion homeostasis, internal modifications and interactions with proteins. For example, melting of RNA base pairs in response to temperature changes can lead to RNA structural transitions (Nikolova and Al-Hashimi 2010), which can be directly linked to translational regulation as a result of start codon accessibility in mRNAs and ribosomal recruitment for translation initiation process (Narberhaus 2010; Kramer and Gregory 2018; Mustoe et al. 2018). Likewise, drug or metabolite sensing of RNA structures are known to effect mRNA stability or translation efficiency in both prokaryotes and eukaryotes (Barrick and Breaker 2007; Cheah et al. 2007; Henkin 2008; Breaker 2018; Warner et al. 2018). In addition to sequence-specific recognition of RNA by RNA binding proteins (RBPs) (Jones et al. 2001; Sanchez de Groot et al. 2019), chemical modification of RNA nucleotides, such as N6-methyladenosine (m6A) modification (Roundtree et al. 2017) provides an added layer of regulation which can significantly impact gene function (Sanchez de Groot et al. 2019; Corley et al. 2020). Recent developments in high throughput sequencing technologies and their coupling with RNA structure-probing approaches now provide a comprehensive platform to map the secondary structure of the whole transcriptome of yeast, plants, metazoans and mammalian cells and functionally relevant RNA motifs (Kertesz et al. 2010; Li et al. 2012; Spitale et al. 2013; Ding et al. 2015; Kwok et al. 2015; McGinnis et al. 2015; Smola et al. 2016; Smola and Weeks 2018). However, our knowledge of these RNA structural motifs in lower eukaryotes, especially in protists, many of which cause deadly human diseases, are limited due to lack of knowledge on their genome-wide abundance of RNA secondary structures and their influence in gene regulatory processes.

*Plasmodium falciparum* is a parasitic protist which is responsible for the deadly disease, malaria (Chen et al. 2000). More than 3 billion people live in areas that are at high risk for malaria transmission (Val et al. 2019). Malaria, a febrile illness, is often characterized by cyclical changes in body temperature (Oakley et al. 2011), which can lead to increased adhesive properties of *P. falciparum* infected red blood cells (iRBCs) (Zhang et al. 2018), a critical contributor to malaria pathology. Also, biophysical studies have recently highlighted temperature dependence of receptor-ligand interactions in *P. falciparum* pathogenesis (Lim et al. 2020). Since temperature fluctuations can greatly affect RNA structures and folding energies, which can ultimately influence RNA stability, translation (Wan et al. 2012; Qi and Frishman 2017) and potentially affect virulence properties of pathogenic organisms (Brower-Sinning et al. 2009; Leach and Cowen 2013; Meyer et al. 2017), it is crucial to get a clearer view of how RNA folding can affect gene function in a lethal disease like malaria. Since the sequencing of the malaria genome, numerous studies have shown that transcription in malaria parasite *P. falciparum* is developmentally regulated (Coulson et al. 2004; Le Roch et al. 2004; Llinas and DeRisi 2004; Hughes et al. 2010). Transcriptome wide studies revealed that the *P. falciparum* asexual stages in human RBCs have a cyclic pattern of steady‐state mRNA expression, with more than 75% of the genes achieving high abundance of mRNAs at only one time‐point of their 48 hours life cycle (Bozdech et al. 2003; Le Roch et al. 2003; Llinas et al. 2006). Additionally, widespread translational delay was reported in malaria (Bunnik et al. 2013). However, gap remains in our current understanding on how this steady state, timely mRNA expression relates to delay in translation, as transcription and translation of malaria genes appear to be tightly linked processes (Caro et al. 2014). Therefore transcriptome-wide control of gene expression and translation remains an active area of exploration in malaria research. In addition, several studies have indicated that post-transcriptional control mechanisms are major means of gene expression regulation in malaria parasites in both sexual and asexual stages. Global ribosome profiling, mechanistic studies on RNA binding proteins (RBPs) (Mair et al. 2006; Sims et al. 2009; Mair et al. 2010; Otto et al. 2010; Caro et al. 2014; Zhang et al. 2014b; Lu et al. 2017; Rios and Lindner 2019) and differential gene expression studies have shown that a wide array of genes, including virulence genes in malaria parasites, are regulated at both post-transcriptional and translational level (Amulic et al. 2009; Amit-Avraham et al. 2015; Chan et al. 2017). RNA-binding proteins (RBPs) are abundant in *P. falciparum* and could particularly influence post-transcriptional genetic control cascades in malaria (Bunnik et al. 2013; Reddy et al. 2015). A recent survey of m6A modifications, which affect translation by resolving mRNA secondary structures (Liu et al. 2015), suggest that these modifications can dynamically calibrate transcriptional and post-transcriptional processes in *P. falciparum* asexual cycle (Baumgarten et al. 2019). Malaria parasites also express antisense RNAs (Militello et al. 2005; Lopez-Barragan et al. 2011; Siegel et al. 2014) that are considered gene regulatory in nature (Gardiner et al. 2000; Epp et al. 2009; Amit-Avraham et al. 2015) and several other noncoding RNAs (Chakrabarti et al. 2007; Mourier et al. 2008; Vembar et al. 2014; Broadbent et al. 2015), many of which fold into characteristic secondary structures (Chakrabarti et al. 2007) that could account for their diverse functional activities. Therefore, like human, yeast and plants (Wan et al. 2011; Rouskin et al. 2014; Wan et al. 2014; Foley et al. 2015), structural contents of RNA could act as a major force of post-transcriptional and translational regulation in malaria. Indeed, early evidence from differences in structural content of large subunit ribosomal RNAs (rRNAs), between asexual and sexual developmental stages, indicated dynamic changes in these rRNAs related to stage-specific ribosome function in *P. falciparum* (Rogers et al. 1996; Hughes et al. 2010).

Given the plethora of observations on the functional importance of the RNA secondary structures in general and RNA-mediated genetic regulation in malaria parasites, we sought to globally monitor changes in RNA structures in distinct asexual developmental stages of *P. falciparum* by combining nuclease mapping with high throughput sequencing. Additionally, we were able to purify biologically relevant antisense RNA regions from total RNA by immunoprecipitation with antibodies that recognize the double-stranded nature of these RNA molecules. Our RNA mapping data not only aligns with known noncoding RNAs that have defined cellular function, it also provides the first genome-wide view of the structural landscape of messenger RNAs (mRNAs) and novel noncoding transcripts which could play important roles in the biology and pathogenesis of malaria. Thus, our findings provide an extensive collection of RNA secondary structures in *P. falciparum* and a valuable new resource for the analysis of RNA structure-function in this organism.

## Results and Discussion

### Transcript assembly and analysis of duplex RNA-seq from *P.falciparum* developmental stages

When studying this pathogen, the primary focus for this project was to look into the RNA secondary structures that were formed in the coding and noncoding segments of transcripts at the genome-wide level. Since conventional RNA interference (RNAi) machinery that produce double-stranded RNAs (dsRNAs) (via siRNA and miRNA mediated pathways) for gene silencing (Carthew and Sontheimer 2009) is absent in malaria pathogen *P. falciparum* (Baum et al. 2009), to avoid any confusion we refer to the double-stranded regions identified in our experiments in *P.falciparum* as ‘duplex RNAs’ or ‘RNA duplexes’, similar to how they are denoted in previous studies (Piao et al. 2017; Lu et al. 2018). To identify RNA duplexes, i.e. secondary structures that are results of inter- and intra-molecular base pairing interactions by deep sequencing, we optimized a protocol (referred here as ‘duplex RNA-seq’) that was originally developed for double-stranded RNA Sequencing (dsRNA-seq) in plants (Zheng et al. 2010; Li et al. 2012; Foley et al. 2015) and obtained base-paired RNAs from the asexual blood stage cultures of malaria parasite *P. falciparum*. Synchronized cultures of *P. falciparum* were collected from the 48 hours asexual RBC cycle and labeled as follows: Early or Ring stage collection: 2-12 hours, Mid or Trophozoite stage collection: 18-26 hours and Late or Schizont stage collection: 30-42 hours post-invasion of parasite in human RBCs. Three samples were collected and pooled for each stage during *P. falciparum* asexual blood stage development (**Figure 1A**). Replicates of these synchronized (>90%) parasite cultures were subjected to RNA isolation by TRIZOL (Invitrogen) method. Total RNA isolated was subjected to two rounds of rRNA depletion using custom made oligos against *P. falciparum* rRNAs (see ‘Materials and Methods’ and **Suppl. Table-1**). These RNA samples were then treated by RNase One (Promega), a single-stranded RNA (ssRNA)-specific endoribonuclease to destroy single-stranded regions of the transcriptome while keeping the base-paired regions intact. Following nuclease treatment, *P. falciparum* duplex RNAs were purified and stage-specific libraries were subjected to RNA sequencing to obtain the first assembly of *P. falciparum* strand-specific transcripts, representing duplex RNA abundance from sequencing data. Two biological replicates of each stage-specific sample were sequenced (see ‘Materials and Methods’ for details) and the sequenced reads obtained from duplex RNA-seq were aligned to the 3D7 reference genome (PlasmoDB, Release 34) using STAR aligner (Dobin et al. 2013) (see ‘Materials and Methods’ for details about the analysis). We used two different Illumina sequencing platforms for two separate rounds of duplex RNA-seq, generating the largest number of raw reads of 67,970,256 and 437,153,310 from HiSeq 2500 and NextSeq500 runs respectively (**Suppl. Table 2**). The read lengths varied from 75bp to 125bp. Correlation analysis between the biological replicates of the structural RNA-seq normalized gene counts (REC) (see Materials and Methods) across the three time point collections of *P. falciparum* reveals high correlation of gene counts between the replicates across these two platforms (**Suppl. Fig 1**). It is worth noting that the duplex RNA-seq measured abundance of base-paired RNA regions among developmental stages and thus, may not be quantitated the same way as total RNA-seq data. Despite this, and differences in sequencing instruments, we see a high correlation of gene counts between biological replicates of the same stage (**Suppl. Fig 1**). Assessing sequencing accuracy, we found on average 83.2% of these total reads showed Phred quality score of Q≥ 30. The distribution of GC content in these reads were between 40-50% which is much higher than average GC content of *P. falciparum* genome (~19%) (**Suppl. Table 2**). This GC-richness of duplex RNA-seq reads suggests that these reads are aligning mostly to the euchromatic coding sequences or functionally important noncoding regions in *P. falciparum* genome which are usually high in GC content (Gardner et al. 2002; Chakrabarti et al. 2007).

**Figure-1:**
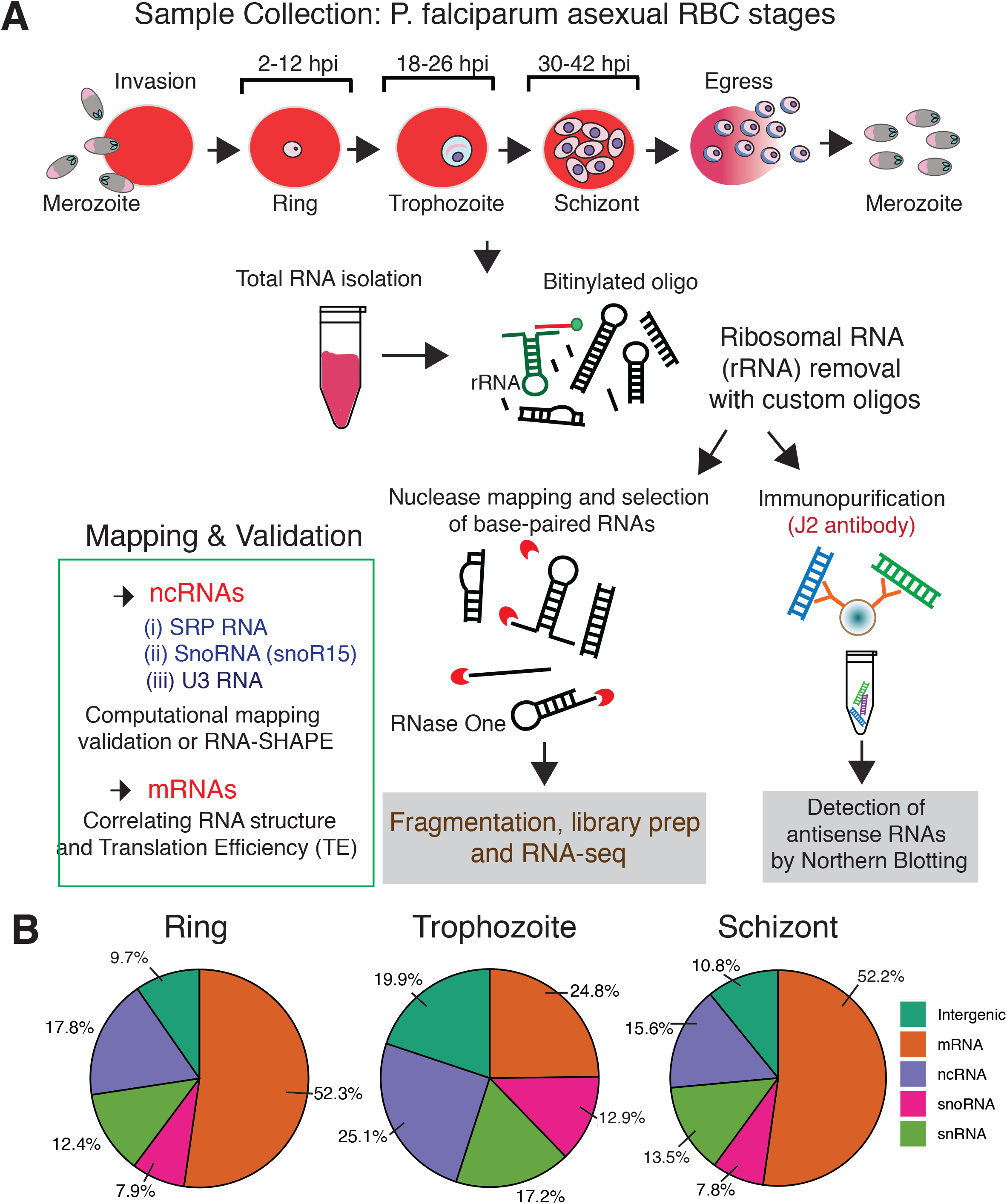
RNA sequencing, analysis pipeline and read distribution profile. A) Total RNA samples collected from three different developmental stages (Ring, Trophozoite and Schizont) of *P. falciparum* in human RBCs were subjected to nuclease-based structure mapping using RNase One, followed by purification of base-paired RNAs and strand-specific RNA sequencing. Differentially expressed regions of the *P. falciparum* genome that show significant accumulation of duplex RNA reads were further analyzed by immunoprecipitation using anti-dsRNA for detection of antisense RNAs. Also, notable structural RNAs that were identified by dsRNA-seq were further validated by mapping to secondary structures following RNA-SHAPE analysis. B) Distribution of paired-end duplex RNA-seq reads that align to different annotated categories of coding and noncoding transcripts in *P.falciparum* genome (PlasmoDB). Only reads mapping uniquely to the genome are considered and reads aligning to known rRNAs and tRNAs are removed from this analysis.

### Overview of duplex RNA distribution and dynamics in *P. falciparum* RBC cycle

As anticipated, a large proportion of global coverage of base-paired RNA reads aligned to genomic regions which produce structural RNAs of known and unknown functions in *P. falciparum* (Chakrabarti et al. 2007; Mourier et al. 2008; Broadbent et al. 2015). Reads mapping to known tRNAs and residual rRNAs were bioinformatically removed before differential expression analysis of duplex RNAs (see ‘Materials and Methods’). Interestingly, a larger proportion of snoRNA, snRNA and known noncoding RNA-specific read accumulations were observed in the trophozoite stage compared to ring and schizont stages of *P. falciparum* (**Figure 1B**). In contrast, ring and schizont stages contained >50% reads that correspond to protein-coding transcripts, representing highly abundant secondary structures within mRNA molecules in these two stages of the *P.falciparum* asexual cycle. On average, the ring and schizont stages have ~23% more “expressed” genes (see expression count calculations in Materials and Methods) containing duplex RNA reads than the trophozoite stage. This data corroborates with stage-specific transcript stabilization that was observed during developmental mRNA dynamics in *P. falciparum* (Painter et al. 2018), probably because RNA stability plays a critical role in mRNA structure driven translational regulation (Mauger et al. 2019).

To better understand *P.falciparum* RNA structurome, we decided to split the coverage by chromosome and determine comparative distribution of the base-paired RNA reads. Thus, we produced a plot of duplex RNA expression from Ring, Trophozoite and Schizont stages on all 14 chromosomes (**Figure 2A**) which revealed that most highly expressed protein-coding genes producing RNA secondary structures in the coding transcripts are residing in chromosomes 6, 11, 13 and 14. Interestingly, many of the antigenic gene families in *P. falciparum* which are involved in host immune evasion and pathogenesis are transcribed from these chromosomes, including multiple copies of *var* and *rif* genes (Gardner et al. 2002). It is noteworthy that the duplex RNA-seq data showed significant enrichment for these mRNAs involved in antigenic variation in *P. falciparum* (see below).

**Figure-2:**
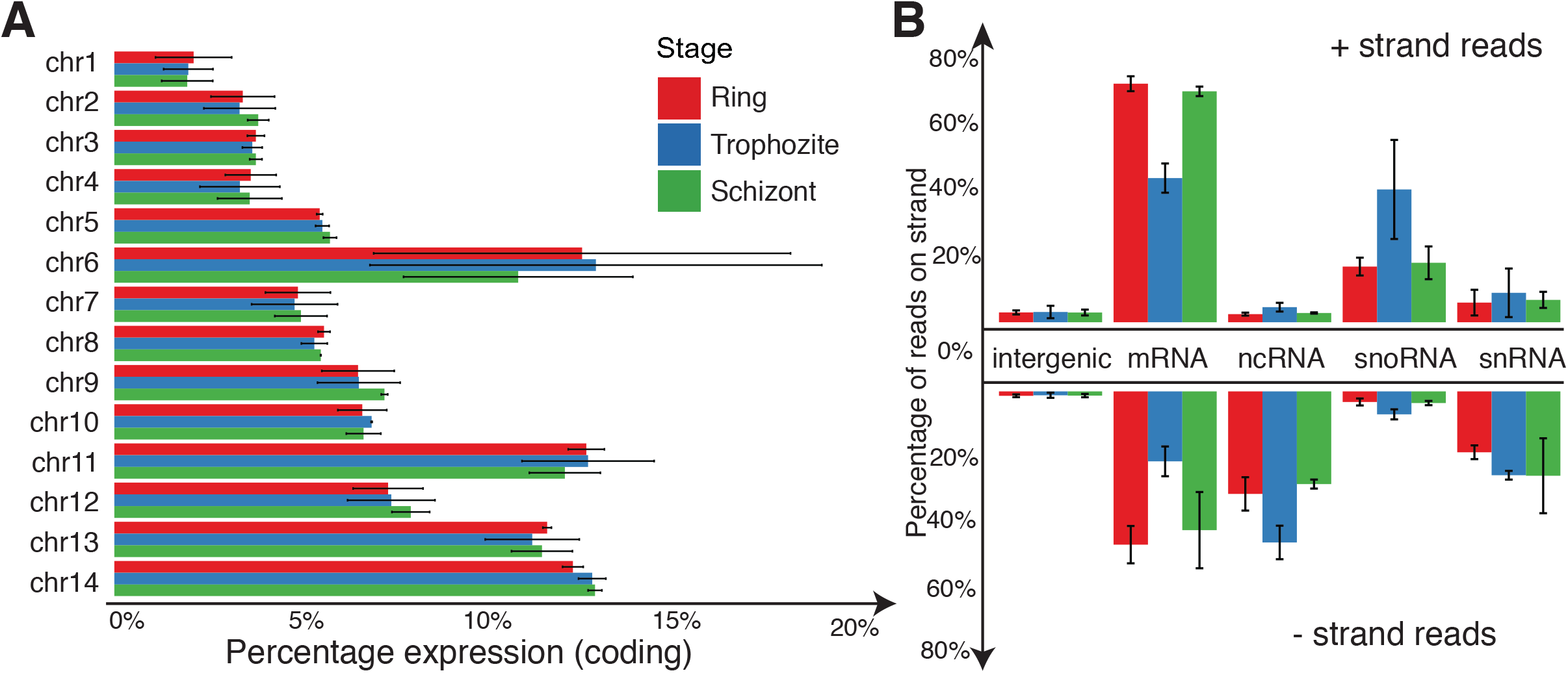
(A) Distribution of uniquely mapped reads across known coding regions on all 14 chromosomes by developmental state; *red:* Ring; *blue:* Trophozoite; *green:* Schizont (B) Distribution of uniquely mapped reads across various annotated coding and non-coding regions of *P.falciparum* genome. The relative percentage of reads from forward stranded reads across various gene categories are shown on top and reads from reverse stranded reads are shown on the bottom. The bars denote average percentage of reads across biological replicates and the error bars show standard deviation.

For known noncoding RNAs, such as snoRNAs, snRNAs, ncRNAs, transcripts from chromosomes 11, 12 and 14 showed highest accumulation of duplex RNA reads and thus, indicate an abundance of structured regions (**Suppl Fig. 2**). On the other hand, duplex RNA enrichment on other chromosomes were very minimal. tRNA and rRNA genes were not included in this analysis. As **Suppl. Fig 2** shows, this observed trend is not simply a result a higher number of non-coding genes on these chromosomes (black dots). The size of non-coding genes also follows a similar pattern as the number of genes, with chromosomes 8 and 11 containing the most area with non-coding genes (~10kB) and chromosome 14, which has the most enrichment for duplex RNA in non-coding regions covers ~6kB (data not shown).

Because the duplex RNA-seq data retained strand information of the molecules, we were able to determine the strand bias for different classes of RNAs. DNA replication and transcription can exert a strand bias on the DNA repair process (Liu et al. 2002; Brachman and Kmiec 2004), particularly transcription levels affecting the degree of bias (Strick and Savery 2017). This in turn could affect an organism’s gene expression program. In *P. falciparum*, strong enrichment of duplex RNA reads transcribed from the forward (plus) strands were observed for mRNAs and snoRNAs from all three stages. Particularly, mRNAs and snoRNAs were ~1.5 fold and ~5 fold enriched respectively for the forward strand compared to reverse (minus) strand across all chromosomes. Expression of mRNAs with secondary structures were higher in ring and schizont stages compared to trophozoite stages, whereas snoRNAs were mostly expressed in the trophozoite stage, irrespective of strand bias (**Figure 2B**). On the contrary, snRNAs and other known noncoding RNAs showed a strong reverse strand bias in the duplex RNA-seq data.

### Genomic distribution of duplex RNA hotspots in *P. falciparum*

The evidence of highly base-paired regions of RNAs, referred hereby as duplex RNA ‘hotspots’ were identified across the coding and noncoding regions of *P. falciparum* genome. From previous studies, it is known that structured regions in the mRNA have the potential to act as regulators in cellular processes, such as RNA stability, turnover and translation in yeast and mammalian cells (Jacobs et al. 2012; Leppek et al. 2018; Mauger et al. 2019). Our duplex RNA-seq data mapped to a total of 3829 protein coding (mRNAs) and non-coding transcripts with relative expression counts (‘REC’) above set thresholds (REC>10) and showed high enrichment (>100 REC) for 969 mRNAs in two or more samples across any of the stages (See Materials and Methods). This suggests that these 3829 coding and noncoding regions could potentially fold into unique, stable secondary structures. A threshold of 10 was set based on empirical observations of peak data being unreliable below this threshold (see Materials and Methods for additional information). In addition, we identified known structural RNAs and novel, structured RNA molecules throughout the intronic and intergenic regions of *P. falciparum* genome.

#### Structural RNAs

Our duplex RNA-seq data is in agreement with known RNA structures in *P. falciparum.* We corroborated our duplex RNA sequencing data with 73 highly base-paired segments of known, conserved structural RNAs (**Suppl. Table 3**). To characterize the constituents of duplex RNA-seq reads, we determined the secondary structure of the *P. falciparum* U3 snoRNA *in vitro* by in-gel RNA-SHAPE analysis using NAI (**Figure 3A**) and mapped our duplex RNA-seq to the U3 secondary structure model. As anticipated, base-paired stem regions originating from intra-molecular interactions in PfU3 RNA corresponded to more duplex RNA-seq reads (represented by peaks in **Figure 3B**) than the unpaired regions (**Figure 3C**). Thus, our analysis revealed with confidence that the majority of our data obtained from duplex RNA-seq are related to biologically relevant structures in the malaria genome. Among all these noncoding RNAs, the highest enrichment of reads (> 333,000 normalized read counts) mapped to the *P. falciparum* Signal Recognition Particle RNA (SRP RNA, PF3D7_1418800) (Chakrabarti et al. 2007), which is a conserved RNA universally required for co-translational protein targeting. The three-dimensional structure of mammalian SRP RNA is already known (Halic et al. 2004) and the secondary structure prediction of *P. falciparum* SRP RNA was previously reported based on comparative genomics and RNA analysis (Chakrabarti et al. 2007), which corroborates strongly with the duplex RNA-seq data (**Figure 3D and 3F**). When stage-specific PfSRP expression was compared to the base-paired transcript abundance in our duplex RNA-seq data (**Figure 3E and 3F**), we observed lower accumulation of SRP RNA domain-specific reads in the ring stage compared to the schizont stage, although expression levels were similar in early ring and schizonts stages as evident from northern analysis. (read accumulations are represented by ‘peaks’ in **Figure 3F** and the missing peak in the ring stage is shown by an arrow in **Figure 3F** with it’s corresponding origin in **Figure 3D)**, This indicated that our duplex RNA-seq was able to capture stage specific RNA structural rearrangements during *P. falciparum* development. Among other highly structured classes of RNA molecules that were identified in *P. falciparum* genome, secondary structure and RNA modification sites for *P. falciparum* C/D box snoRNA 15 (PF3D7_1222200) was previously established by *in vivo* DMS probing (Chakrabarti et al. 2007). Our duplex RNA-seq accurately mapped to those structured regions of *P. falciparum* snoR15 (**Suppl. Figure 3**), suggesting that we are able to reliably capture intramolecular base-pairing interactions by duplex RNA seq and afforded the first genome-wide view of the structural landscape of *P. falciparum* RNAs, many of which are involved in conserved biological functions. In addition, RNAs of Unknown Function (RUFs) (Chakrabarti et al. 2007), RUF1 (PF3D7_1328700) and RUF 6 (PF3D7_0711800, PF3D7_0712100, PF3D7_0712700), all expressed from chromosome 7, showed enrichment of base-paired RNAs.

**Figure-3:**
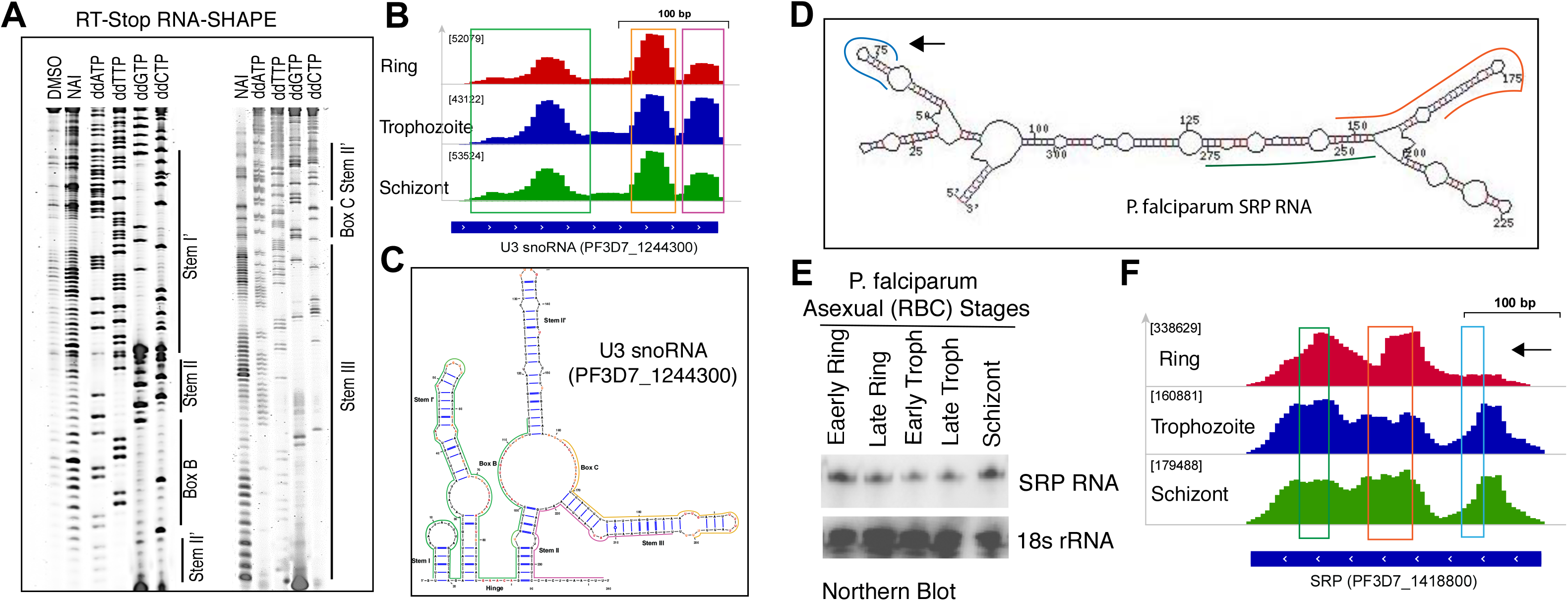
Mapping and validation of duplex RNA-seq data with known RNA secondary structures of noncoding RNAs: U3 RNA (A-C) and SRP RNA (D-F).(A) Two Cy5 labeled denaturing gels for U3 snoRNA domains: Stem I’, Hinge, Box B, Stem II’, Stem II’, Box C and Stem III (Corresponding RNA secondary structure data in panel C). (B) IGV screenshot of the U3 RNA peaks representing duplex RNA-seq read enrichments in three different P. falciparum RBC stages. (C) Secondary structure of the Pf U3 snoRNA based on in-gel SHAPE modification of mix stage RNA using NAI (shown in panel A). Unpaired RNA nucleotides are shown in red and orange color based on their reactivities, where high reactivity is signified by red color and low reactivity by orange with the SHAPE reagent. Base-paired RNA regions identified from duplex RNA-seq in B are indicated by green, yellow and magenta color skirting (also shown by matching colored bars at the bottom of the peaks/reads identified in duplex RNA-seq). (D) Schematic representation of the P. falciparum SRP RNA secondary structure model. Mapped reads from duplex RNA-seq are shown in skirted colored lines along the structure model. Corresponding peaks of the mapped read regions in P. falciparum genome are shown by three color matching boxed regions in F. Due to shorter insert sizes in 3 replicates of the data (**Suppl. Table 2**), the peaks are seen more distinctly and at a higher resolution in these replicates and thus, we selected to display the coverage in these replicates. The remaining three replicates also show the same structures but at a lower resolution. (E) Northern blot analysis of the total RNA from five different time point collections during asexual RBC cycle. (F) Black arrow showing a peak missing from the ring stage samples (also shown by an arrow in structure model in D), indicating the ability of the duplex RNA-seq to detect stage specific RNA structural rearrangements during *P. falciparum* development.

#### Coding Sequence (CDS) of mRNA

Our duplex RNA-seq readout allowed global analysis of RNA secondary structures in the coding regions of *P. falciparum* mRNAs. Recent reports suggest that highly expressed mRNAs usually have extensively structured coding sequence (CDS) (Wan et al. 2014; Mauger et al. 2019), which could positively correlate with transcript-specific translation efficiency by increasing the mRNA half-life (Mauger et al. 2019) and the ribosome (Beaudoin et al. 2018) or m6A modifications (Mao et al. 2019) in CDS or UTRs could be the major remodeler of the mRNA structures for translation. In malaria parasites, mRNA translation facilitates developmental stage transitions (Cui et al. 2015; Painter et al. 2018), suggesting that malaria disease progression is dependent on stage-specific protein output. Additionally, *in vivo* measurements of mRNA cellular dynamics by biosynthetic labeling recaptured the ‘just-in-time’ transcriptional program of *P. falciparum* mRNAs with variable degrees of mRNA stabilization profile (Painter et al. 2018), which could be due to differential presence of structure between these mRNA populations. Therefore, it is plausible that a great proportion of post-transcriptional gene regulatory program is driven by mRNA secondary structures in this parasite. We found that nearly 71.4% of the protein coding genes in *P. falciparum* aligns to duplex RNA-seq reads exceeding our set threshold (REC>10), and as a result, define its secondary structure content. Out of the uniquely “expressed” 3773 mRNA regions containing secondary structures, approximately a quarter (26.1%) show high cumulative expression (>100 REC) of transcripts with structured regions in at least 1 stage. Based on the average expression seen across all the dsRNA-Seq time points, we conclude that approximately 71% (3773) of all mRNAs make secondary structures in at least one of the sequenced stages, with expression above set thresholds. When comparing the distribution of expressed genes (REC) between the stages, we found significant difference among Ring and Trophozoite or Trophozoite and Schizont stages (**Suppl. Figure 4**) (two-sided Wilcoxon test, p < 0.001). However, no significant difference was observed between the distributions of the expressed genes of Ring and Schizont stages in the *P. falciparum* RBC cycle.

In order to study the correlation between the gene expression of protein coding transcripts in *P. falciparum* total RNA-seq (Otto et al. 2010) and the duplex RNA-seq data, we normalized each dataset separately using DESeq2 (Love et al. 2014) and plotted the average normalized gene expression counts for each stage across the total RNA-seq and duplex RNA-seq datasets. As expected, the protein coding genes have more expression in the mRNA-seq dataset compared to the duplex RNA-seq as the latter represents a subset of the former, containing expressed structure contents of the protein coding genes (**Figure 4A, Suppl. Figure 5A** and **5B**). The overall degree of correlation between the two datasets is quite high (R^2^ values are 0.66, 0.65, 0.70 across the three stages). Few genes show higher expression in duplex RNA-seq compared to mRNA-seq and few of these are highlighted in the figures.

**Figure-4:**
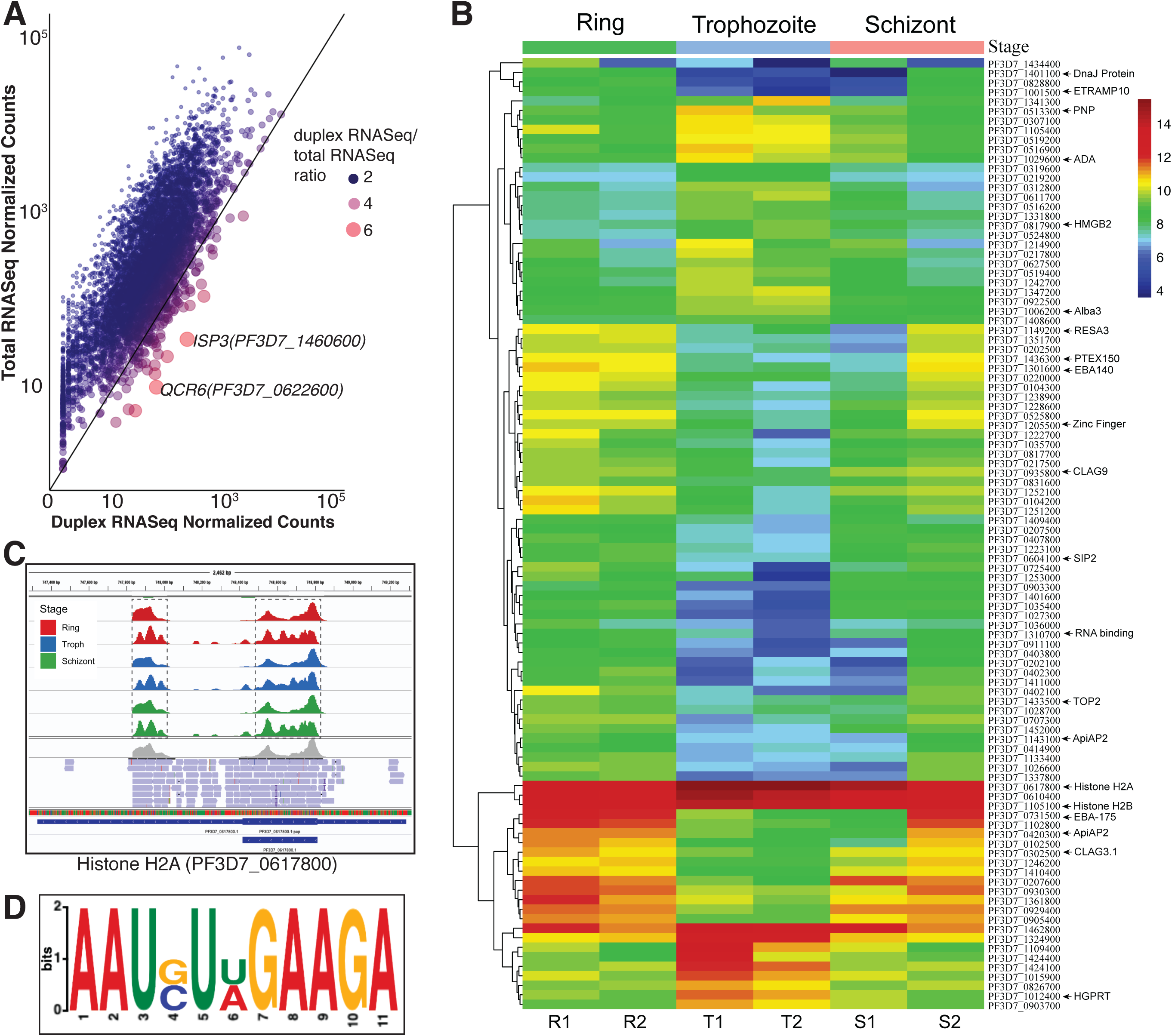
(A) Gene expression of duplex RNA-seq as compared to *P. falciparum* total RNA-seq data (Otto et al. 2010). Representative scatter plot showing correlation between normalized gene counts from total RNA-seq and duplex RNA-seq datasets for Ring stage. The other stages are shown in **Suppl. Fig. 5A and 5B**. A pseudocount of 1.0 is added to the normalized counts for both duplex and total RNA-seq data prior to plotting. The axes are in logarithmic scale. The color and size of the points show the ratio of the duplex RNA-seq normalized counts by the total RNA-seq normalized counts. Larger red points represent genes with higher expression in duplex RNA-seq compared to total RNA-seq as shown in the legend. (B) Heatmap of the 100 more expressed differentially expressed genes across the three developmental stages. The normalized gene counts from DESeq2 are log transformed in the heatmap. The highlighted genes on the right are referenced across the text. (C) Screen shot of duplex RNA enrichments for all six samples for Histone H2A visualized by IGV browser. The gene contains two sets of peaks in the duplex RNA-seq data, highlighted by dashed boxes. The second replicates show a higher resolution of peaks due to shorter insert sizes (as discussed before) (D) A highly represented sequence-structure motif identified from duplex RNA reads of *P. falciparum* mRNAs.

To understand the mRNA structure dynamics during *P. falciparum* asexual stage transitions in human RBCs, we characterized differential expression of structured contents within mRNA coding sequences (CDS), which is depicted as a heatmap in **Figure 4B**. Results from the top 100 most expressed mRNAs with secondary structures (out of all the differentially expressed ones) during the 48 hours intra-erythrocytic developmental cycle (IDC) of this parasite suggest that highly folded RNA coding regions can potentially influence stability and translational outputs of individual transcripts in malaria gene expression programs. Among the highly expressed base-paired mRNA molecules in *P.falciparum*, histone mRNAs (Histone H2A (PF3D7_0617800), H2B (PF3D7_1105100) and H3 (PF3D7_0610400)) showed highest accumulation (within top 1%) of reads by duplex RNA-seq. Histone mRNAs are one of the most extensively validated functional mRNA structures present not only in metazoans, but also in many different protozoa, including alveolates (e.g. *P. falciparum)* (Davila Lopez and Samuelsson 2008). The histone mRNA forms a stable stem-loop structure at the 3’ UTR structure, which is involved in nucleocytoplasmic transport and regulation of translation efficiency (Zhang et al. 2014a). In addition to strong enrichments of duplex RNA reads at the 3’ UTR of *P. falciparum* histone mRNAs, read abundance was also observed in the CDS, as shown for histone H2A (**Figure 4C**). In addition, the list of highly expressed duplex RNA containing transcripts in *P. falciparum* included mRNAs of chaperon proteins *HSP70* and *HSP90*, which are also known to be highly structured (Ahmed and Duncan 2004; Sanchez de Groot et al. 2019). These findings substantiated that our duplex RNA-seq interrogated the desired regions of mRNAs on a genome-wide scale, which are highly enriched in base-pairing interactions.

One emerging theme about *P. falciparum* gene regulation is the central role played by DNA and RNA binding proteins in transcriptional and post-transcriptional control of gene expression (Vembar et al. 2016; Painter et al. 2018; Rios and Lindner 2019). For example, a plant-like transcription factor family consisting of 27-members, known as Apicomplexan AP2 (*ApiAP2*) protein family, expressed in different stages during malaria parasite development, is critically involved in parasite development and differentiation processes (Balaji et al. 2005; Painter et al. 2018). Expression of two members of this gene family, PF3D7_0420300 and PF3D7_1143100 (**Figure 4B**), were among the top 100 gene candidates that produced high levels of differentially expressed duplex RNAs in their coding regions and thus considered in our study as prominent ‘hotspot’ candidates. Among those, *ApiAP2* (PF3D7_0420300) expressing protein is known to functionally interact with ACACACAT DNA motif for *P. falciparum* gene expression regulation (Toenhake et al. 2018) and its expression is correlated with drug resistance transporter, *pfCRT*, indicating potential role in resistance evolution to chemotherapeutic molecules in malaria (Siwo et al. 2015). Similarly, for RNA mediated post transcriptional gene regulation, RNA binding proteins (RBPs) in *P. falciparum* plays pivotal role in translational regulation (Bunnik et al. 2016; Rios and Lindner 2019). Our duplex RNA-seq data revealed strong accumulation of base-paired reads mapping to DNA/RNA binding factor *PfAlba3* (PF3D7_1006200) (**Figure 4C**). *PfAlba3* protein interacts with *P. falciparum* histone deacetylase, *Sir2A* (Goyal et al. 2012), associates with polysome fractions during the *P. falciparum* IDC (Bunnik et al. 2016) and therefore, may play important regulatory roles in transcription as well as in translation. In addition to mRNAs that produce regulatory proteins, transcripts encoding antigens and adhesion molecules involved in host-parasite interaction in *P. falciparum* are the most highly base-paired classes of RNA molecules, including Erythrocyte binding antigen 175 (*EBA-175*) (PF3D7_0731500), which is involved in erythrocyte invasion (Mamillapalli et al. 2006), *SERA5* (PF3D7_0207600), important for parasite egress (Collins et al. 2017) and ring-infected erythrocyte surface antigen, *RESA-1* (PF3D7_0102200) and *RESA-3* (PF3D7_1149200) (Favaloro et al. 1986), which may prevent *P. falciparum* infected erythrocytes from reinfection through interactions with spectrin family proteins (Pei et al. 2007).

Genetic control of cellular metabolism often involves sensing of mRNA secondary structures by nutrients and metabolites or interactions with RBPs (Sudarsan et al. 2003; Clingman and Ryder 2013). These mRNA structures are widespread in prokaryotes and also identified in handful of eukaryotes. In prokaryotes, these are referred to as RNA genetic switches which are typically located in noncoding regions of mRNAs (such as UTR regions) of many metabolic enzymes and can modulate gene expression. In eukaryotes, metabolic enzymes can interact with mRNAs to affect mRNA stability or translation efficiency (Clingman and Ryder 2013). We have identified base-paired mRNA elements in the coding region of *P. falciparum* metabolic enzymes, with most enrichment of duplex RNAs in the Hypoxanthine-guanine phosphoribosyltransferase (*HGPRT*) (PF3D7_1012400) and Adenosylhomocysteinase (*SAHH*) (PF3D7_0520900). Since most parasitic protozoa do not have the *de novo* purine nucleotide biosynthesis pathway and rely mostly on the salvage pathway for purine metabolism where hypoxanthine is the key precursor, *HGPRT* in this pathway makes a promising drug target for *P. falciparum* infection (Cheviet et al. 2019). However, the cellular interactions and importance of these novel mRNA structural elements in the coding regions of metabolic enzymes remain to be analyzed.

The importance of sequence and structural motifs in mRNAs are vastly appreciated in studying post-transcriptional regulatory processes (Rabani et al. 2008; Dominguez et al. 2018). In order to determine if there were any sequence motifs located within the structured regions of mRNAs across highly expressed pool of transcripts, we obtained sequences from 50 highly expressed mRNAs representing a defined structured region (structured regions represented by peaks shown in **Figure 4C**). We then used the pattern search program, Improbizer (Ao et al. 2004) that searched through duplex RNA read sequences and looked for reoccurring short sequence motifs and found six motifs which could be potential regulatory region or an RBP interaction site on mRNA structures. The highest scoring motif sequence is shown as sequence logo in **Figure 4D**. In summary, the overall abundance (**Figure 1B, 2B**) and dynamic changes (**Figures 4A, 4B**) in mRNA structures during *P. falciparum* blood stage development indicate that the extensive stage-specific RNA folding events can be potentially linked to fine-tuning of protein synthesis in malaria gene expression program.

#### Noncoding regions of mRNAs - 5’ and 3’ UTRs

To investigate the global changes in *P. falciparum* mRNA structure during asexual blood stage developments, especially in the untranslated regions (UTRs), we analyzed top 20% differentially structured regions over development in the UTR regions. Since the UTR lengths have a very broad range in the *P. falciparum* genome, the UTRs were binned into 50 and 500 nucleotides (5’ and 3’ UTRs: defined by (Chappell et al. 2020); CDS: 100 nt downstream of the start codon (for 5’UTR) and 100 nt upstream of the stop codon (for 3’UTR), calculated average expression from ring, trophozoite and schizont stages (see ‘Materials and Methods’ for details about the analysis). **Figure 5A** and **5B** show the distribution of normalized expression ratios of the duplex RNA-seq reads at the 5’ and 3’ gene ends for the top 20% genes with structures in the UTRs respectively. The ratio is calculated against the maximum expression of the structures in the region being investigated in order to study the relative enrichment (See Methods). For example, the Schizont structures at the UTRs and splice junctions show lower ratios and thus, lower relative enrichment than the Ring and Schizont stages (**Figure 5A-C**) (t test, p < 0.001). Having secondary structure near areas of translation initiation sites or coding regions have been seen to play a role in regulating translation efficiency. For non-protein coding regions, i. e. 5’ UTRs and 3’ UTRs of mRNAs molecules, there is a significant enrichment in longer UTRs in the Ring and Schizont stages compared to the Trophozite stage (**Suppl. Fig 6**) (Wilcoxon test p-value <0.01).

**Figure-5:**
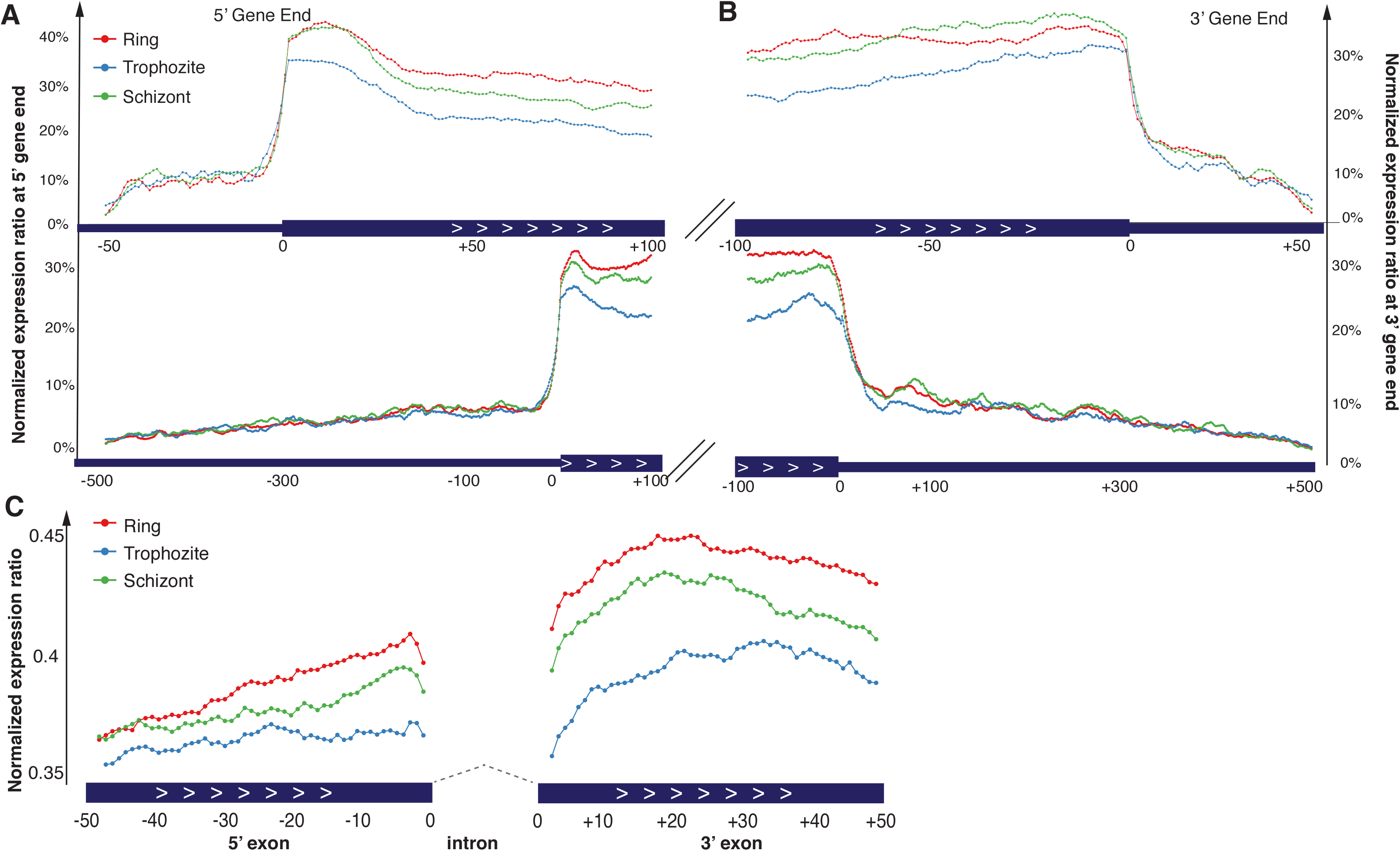
Distribution of duplex RNA-seq data across the top 20% expressed (A) 5’ UTRs, (B) 3’UTRs and (C) splice junctions. The y-axis shows the normalized expression ratio, which is normalized against the highest duplex RNA-seq expression in the region being investigated. This ratio represents the relative distribution of enrichment of base-paired RNAs (See Methods). For the UTRs, all UTRs shorter than 500bp (the median UTR length) are binned into 50bp bins and all UTRs longer than 500bp are binned into 500bp bins, shown by the top and bottom panels in (A) and (B). We focus on 100bp of the coding regions downstream of the start codon and upstream of the stop codon for the 5’UTR and 3’UTR respectively. (C) Coding regions of 50bp flanking the splice junction is plotted. The values for the replicates for each stage are averaged for plotting.

### Antisense RNAs

Antisense RNA transcripts, originated from the opposite strand to a protein or RNA-coding strand, have been ascribed roles in gene expression regulation by degradation of the corresponding sense transcripts or by silencing genes at the chromatin level (Faghihi and Wahlestedt 2009). Therefore, pervasive antisense transcription in *P. falciparum* fueled the speculation that these RNAs regulate malaria gene expression programs either at the epigenetic or post-transcriptional level (Gunasekera et al. 2004; Lopez-Barragan et al. 2011; Jiang et al. 2013; Siegel et al. 2014). At the transcriptome level, widespread antisense transcription in *P. falciparum* corresponded to transcription initiation in an antisense orientation to coding genes(Adjalley et al. 2016), transcription start site-associated RNAs (TSS-RNAs) expression (Chappell et al. 2020) as well as antisense transcription in the context of pre-mRNA splicing events(Sorber et al. 2011). At the gene level, natural antisense transcripts (NATs), longer than 200 nt and acting as antisense long RNAs can potentially regulate *P. falciparum* virulence gene expression (Epp et al. 2009; Amit-Avraham et al. 2015), however, the origin and the mechanism of action of these long antisense RNAs transcribed from either ‘silent’ or ‘active’ var genes, remains unclear (Ralph et al. 2005). To investigate if our duplex RNA reads constitute potential antisense transcripts in *P.falciparum* genome, we counted the antisense reads for all coding and non-coding transcripts and normalized these counts by depth of sequencing (REC). The same threshold described earlier (REC>10.0 in at least two samples across any of the three RBC stages of *P. falciparum*) was applied to the antisense data as this was the empirically set threshold for more reliable duplex RNAs. Since antisense reads are in low abundance (after applying the above stringent cutoff), we found a total of 159 genes (154 mRNAs and 5 non-coding RNAs) with antisense reads above set thresholds. To better understand the relationship between gene expression and the abundance of duplex antisense reads, we looked at the average expression of the top 50 mRNA genes with antisense expression (**Figure 6A)**. As shown in the **Figure 6A**, the gene expression from mRNA-seq and duplex antisense expression are not correlated but show an approximately negative relationship (Pearson correlation R^2^ = −0.11). This means that for certain genes, such as *PF3D7_1425200, PF3D7_0112800* etc, the lower mRNA expression might be explained by antisense transcriptional regulation, although this remains to be validated by functional assays. We saw a few interesting types of antisense expression, especially in the 3’ UTR regions (36.5% antisense structures were found in the 3’UTR). **Figure 6B** represents the three types of antisense peaks seen in the 3’ UTRs (i) antisense peak without any overlapping genes (ii) antisense peak with overlapping UTR of another protein coding gene, and (iii) antisense peak from overlapping non-coding gene(s). An example of (iii) can be seen in **Figure 6C**. To validate antisense RNA expression, we immunoprecipitated rRNA depleted *P. falciparum* total RNA with dsRNA specific J2 monoclonal antibody (SCICONS) and probed this RNA with sense and antisense RNA specific complementary oligonucleotides. In this process we confirmed expression of ten (10) sense/antisense RNA pairs from asynchronous, mix stages of RBC cycle. Representative northern blots validating presence of antisense RNAs opposite three (3) genes are shown in **Figure 6D**. Although our deep-sequencing data and Northern blots correspond to the overlapping regions of the RNAs, it remains unclear whether mRNA stability was affected by the expression of these antisense RNAs or their effect in resolving secondary structures of translatable mRNAs in *P. falciparum*.

**Figure-6:**
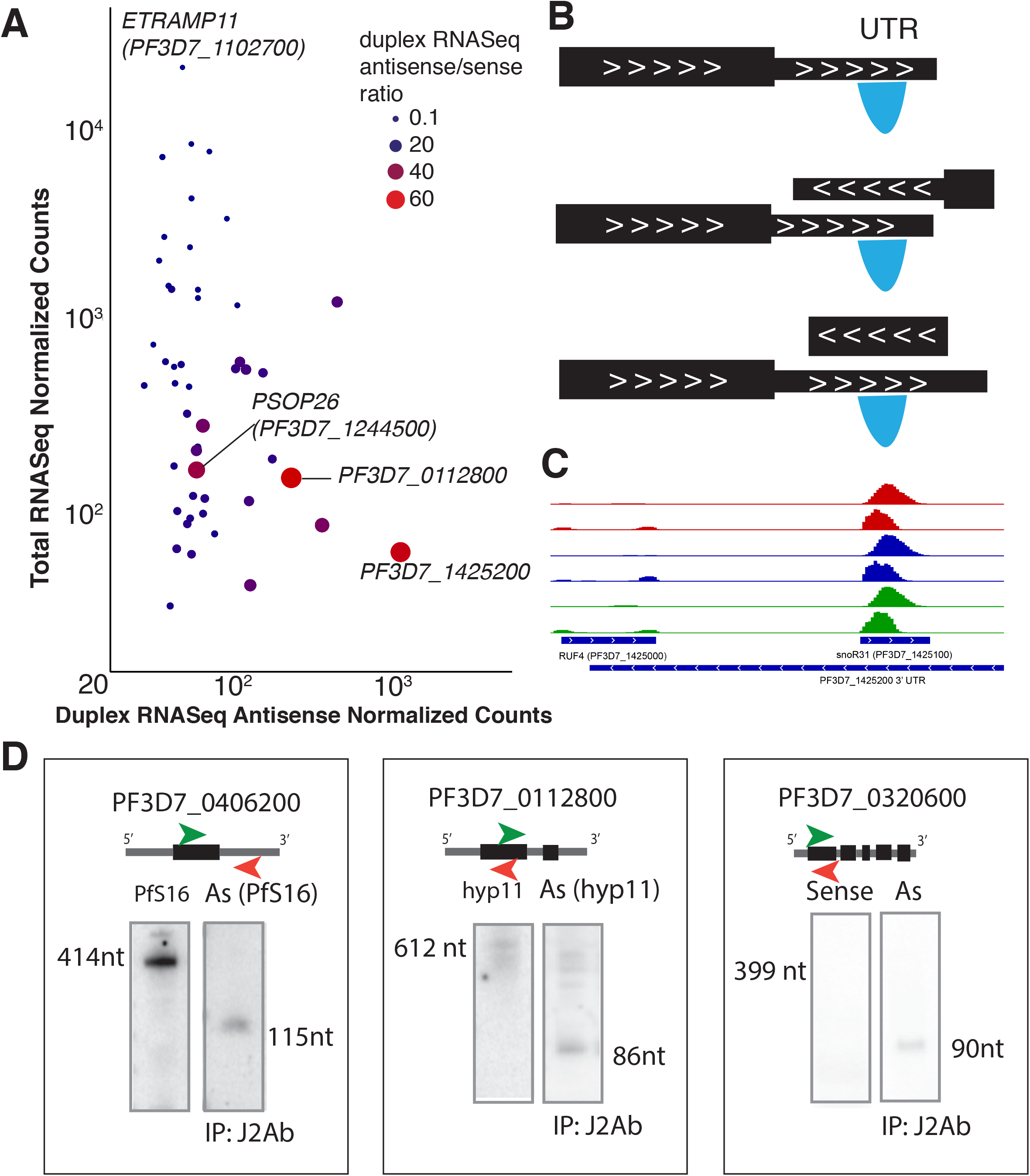
Antisense expression in duplex RNA-seq (A) Antisense duplex RNA-seq counts for the top 50 antisense genes plotted against mRNA-seq counts for Ring Stage. The other stages are shown in Suppl. Fig 4C-D. The size and color of the points indicate the ratio of the duplex RNA-seq antisense to sense counts. Larger red points indicate genes that have more antisense signal than sense signal. Few selected genes with very high and very low ratios are highlighted. A pseudocount of one is added to the normalized counts and axes are in logarithmic scale. (B) A schematic indicating the three types of antisense peaks seen in the duplex RNA-seq data in the 3’ UTRs of the genes (i) antisense peak without any overlapping genes (ii) antisense peak with overlapping UTR of another protein coding gene (iii) antisense peak from overlapping non-coding gene(s). (C) An example gene with high antisense peaks originating from non-coding snoRNA and ncRNA (snoR31 and RUF4) across all three stages is shown. (D) Antisense transcripts detected by northern blotting in the J2 antibody immunoprecipitated samples from mix (asynchronous) stages *P. falciparum* total RNA. Left blot (in all panels) shows sense gene expression and the right blot shows antisense expression from immunoprecipitated samples.

### Genome-wide relationship between RNA secondary structures and biological processes in *P. falciparum*

Next we investigated genome-wide relationships between mRNA structures and biological functions of the encoded proteins. For this purpose, we used differential expression data of secondary structures from the three *P. falciparum* developmental stages within RBCs, i.e. ring, trophozoite and schizont stages (**Figure 4B**). Differential expression of structured mRNA can be measured through the read count in the different stages where a higher read count in one stage versus the other is representative of stage-specific differential accumulation. A GO enrichment analysis of the differentially structured transcripts shows that the genes are enriched for molecular functions that are highly relevant for the developmental stages. These processes include host cell surface binding (Q < 1 × 10^−3^), DNA/RNA binding, which includes mRNA binding (GO:0003729) and sequence-specific DNA binding (GO:0043565) and multiple GO terms that belong to processes involved in enzyme activity. Among these genes, are transcription factors like Plant-like transcription factor *ApiAP2* and *ALBA3* (PF3D7_1006200) ApiAP2 protein *SIP2* (0604100) (P<0.05 Ring vs Trophozoite stage). In addition to most significantly enriched molecular functions corresponding to proteins involved transcription and RNA processing, our duplex RNA-seq data showed GO enrichment for processes, such as red blood cell invasion and host immunity (for example, *EBA-181* and EBA-175, P<0.05, Ring vs Trophozoite stage). Thus, systematic analysis of mRNA secondary structures in gene expression and molecular function in *P. falciparum* identified candidates that are substantially enriched for functions relevant to malaria biology and pathogenesis.

### Analysis of mRNA structure remodeling and translational control

Dynamics of mRNA unwinding can significantly affect the rate of translation in eukaryotic systems. Previous studies have suggested that stable secondary structures in the 5’UTR or coding regions (CDS) of mRNAs can lead to ribosomal pausing our translational arrests (Mao et al. 2014; Rodnina 2016). Given the critical role mRNA structures can play in translation elongation, we next sought to determine the correlation between structural contents of coding regions and translation efficiency in *P. falciparum*. Using existing ribosome profiling data sets (Caro et al. 2014), we used the ribosomal footprint RPKM as a representation of translation efficiency in *P. falciparum* and compared it to the duplex RNA-seq structure levels. While looking at the top 50% structure-containing transcripts in the *P.falciparum* genome, our developmental stage-specific data, represented by the expression of the genes in the duplex RNA-seq datasets and the translational efficiency of these genes revealed an overall positive correlation for most of the mRNA coding regions (R = 0.49, 0.51, 0.53 for Ring, Trophozoite and Schizont p<0.001) (**Figure 7**). However, how individual transcript-specific differences in RNA structures relate to translation efficiency still needs to be determined for *P. falciparum* mRNA translation.

**Figure-7:**
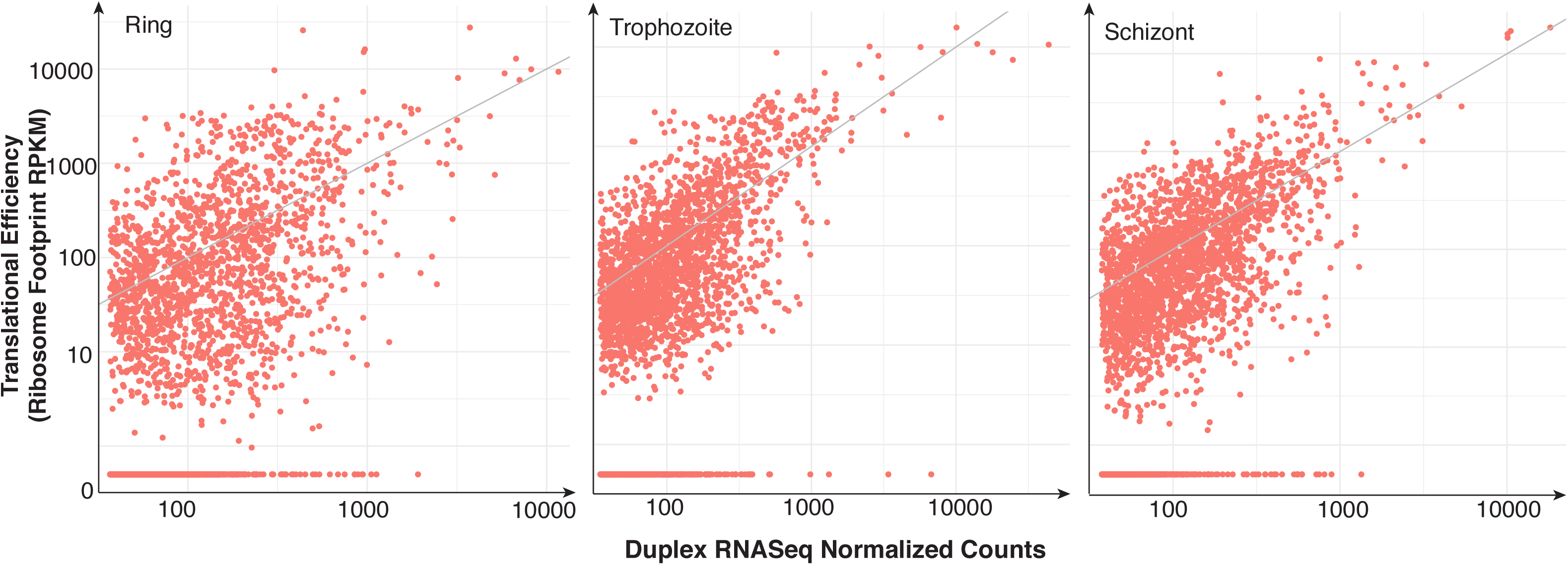
Translation efficiency correlation with duplex RNA-seq structures. Scatter plot of normalized duplex RNA-seq counts plotted against ribosome footprint RPKM data (translation efficiency) from Caro et. al. 2014 for the (A) Ring stage (B) Trophozoite and (C) Schizont stages. The top 50% protein coding genes with structures are plotted. The Pearson correlation coefficient for the three stages are 0.49, 0.51 and 0.53 respectively (p < 0.001).

## Conclusions

Our analysis of *P. falciparum* RNA structurome elucidated stage-specific RNA structural changes and therefore has a potential to yield significant insights into structure-function relationships in post-transcriptional control. Our comparative analysis with other eukaryotic RNA structuromes revealed that *P. falciparum* contains substantially high secondary structures (~71.4%) in protein coding regions compared to human, yeast and metazoan species. Given that *P. falciparum* genome has the highest AT content (~80%) of any organism sequenced so far, it is striking that *P. falciparum* transcriptome contains substantial stable, dynamic secondary structures compared to other organisms with higher GC content. Three membrane layers of host and the parasite have posed a challenge for in-cell probing of malaria structure by high-throughput methods. By overcoming this using a robust *in vitro* assay, we show that reliable detection of structural contents in malaria transcriptome is feasible by duplex RNA-seq. Many of these structural regions in coding sequences are phylogenetically conserved at the sequence level, providing an opportunity to study RNA-structure-dependent functions in malaria animal models. Detailed interrogation of how these structures in malaria coding regions relate to function should provide mechanistic details for developing future therapeutics affecting mRNA structure and stability differently, depending on how they inhibit malaria translation.

## Materials and Methods

### Plasmodium falciparum Culture, Synchronization and Treatments

*P. falciparum* laboratory adapted parasite line 3D7 was cultured at 2% hematocrit in purified human erythrocytes/RBCs using standard culturing techniques (Trager and Jensen 1976; Duffy and Avery 2018) using RPMI 1640 medium (supplemented with 37.5 mM HEPES, 7 mM D-glucose, 6 mM NaOH, 25 μg of gentamicin sulfate/ml, 2 mM L-glutamine, and 10% Albumax II (Life Technologies) at a pH of 7.2 in a 3% carbon dioxide environment. The asexual blood stage parasite density was calculated routinely using thick blood films. Baseline RBC count was used to calculate the parasitaemia (parasites/μl). P.falciparum Infected Red Blood Cells (iRBCs) were treated with a pre-warm aliquot of 5% D-sorbitol in RPMI at 37°C to synchronize parasite cultures in order to isolate homogeneous stages of parasites. Multiple sorbitol treatments were made to achieve >90% synchrony of parasites. Parasites were collected from three different developmental stages, Ring, Trophozoite and Schizont, after stringent synchronizations.

### RNA extraction, rRNA depletion and nuclease mapping

*P. falciparum* parasites were isolated after saponin lysis, washed with incomplete medium (without Albumax) and RNA from each stage were isolated by Trizol (Thermofisher) method followed by Turbo DNAse (Thermofisher) treatment. Total RNA was then subjected to two rounds of rRNA depletion with custom biotinylated oligos designed against *P.falciparum* rRNAs (**Suppl. Table 1**). Briefly, 500 pmol of biotinylated capture probes were used in binding buffer containing 10 mM Tris-Cl pH 7.4, 3 mM MgCl_2_, 300 mM NaCl and 0.5% Igepal-CA 630. Hybridization reaction was prepared on ice in presence of 50 ug of total RNA in each tube and 2ul of RNase inhibitor. Oligo probe and total RNA mixture were heated at 70 °C for 5 min followed by gradually reducing the heat until the oligonucleotides have reached room temperature. Finally, rRNA bound oligos were separated by Streptavidin conjugated magnetic Dynabeads M-280 (Invitrogen) followed by ethanol precipitation of rRNA depleted total RNAs. Next, RNA samples were treated with a single-strand specific ribonuclease, RNase One (Promega) (5 units per ug of RNA) according to manufacturer’s instructions. After nuclease treatment, each RNA sample was fragmented with Fragmentation Reagents (Ambion) and then end corrected with T4PNK (Thermofisher), phenol-chloroform extracted and ethanol precipitated before using them for RNA-seq library preparation.

### Library preparation and high-throughput sequencing of duplex RNAs (Duplex RNA-seq)

*P. falciparum* duplex RNA sequencing libraries were prepared from three developmental stages: Ring, Trophozoite and Schizont (see results section for details). Two independent sets of libraries were prepared (2 replicates for each developmental stage). Sequencing library was constructed using the Small RNA Sequencing Kit v3 (Illumina, San Diego, CA) as per manufacturer’s instructions. Library quality was checked by Bioanalyzer and then pair-end sequencing was done on two different Illumina high-throughput sequencing platforms, HiSeq2500 for RNA-seq-1 (Otogenetics Corp., Atlanta, GA) and NextSeq500 for RNA-seq-2 (ACGT Inc, Germantown, MD) (**Table-1**).

### Bioinformatics analysis of duplex RNA-seq data

The paired end reads from the duplex RNA-Seq data were first trimmed to remove any adapter sequences using fastp v0.20.0 (Chen et al. 2018). STAR v2.5.2 (Dobin et al. 2013) was used to align the trimmed data to the *P. falciparum genome* (v34) with the parameters --genomeSAindexNbases 11 and --alignIntronMax 500. The maximum splice junction length was set to a conservative 500bp as a larger value of this parameter yielded many false splice junctions and thus, noisy structural peaks. This is mainly due to the short insert sizes in the libraries (**Suppl. Table 2**) and thus, relatively shorter length of the trimmed reads post adapter trimming. A custom filter was then used to remove low mapping quality (mapping quality =0) and secondary or supplementary alignments. Finally, featurecounts v2.0.0 (Liao et al. 2014) was used to perform gene counting using the latest gene annotations from (Chappell et al. 2020). Reads that mapped to mitochondrial or apicoplast transcripts were removed from this gene expression analysis. Normalization of gene counts (as seen in Fig. 4) and differential gene expression was performed using DESeq2 v1.26.0 in R v3.6.2. Expression count calculations to estimate the expressed genes are described below.

### Expression count calculations to flag potential stable secondary structures

The raw counts from ‘featurecounts’ including all known coding and non-coding genes were normalized by the relative sequencing depth based on the number of mapped reads to calculate the relative expression counts (REC). Only genes with ‘REC’ values greater than 10.0 across 2 or more samples were considered for this analysis. This is so that outlier noise in single samples will be removed from the analysis. Finally, from the filtered genes, all genes with ‘REC’ values >10.0 filter are considered genes with potential stable structures for each sample. A threshold of 10.0 was selected based on manual inspection of the data in Integrative Genomics Viewer (IGV) (Robinson et al. 2011) as genes with lower values were typically non-specific noise. We did not normalize data by gene length (similar to the popular RPKM i.e. reads per kilobase per million values used in total RNA-seq) as structural data is not as quantitative and does not necessarily contain reads across the entire gene. A longer gene might contain the same number of reads as a similarly expressed shorter gene due to structural similarities and thus, using RPKM would penalize the longer genes. We use the ‘REC’ values to get a global picture about the genes that form duplex RNA structures across the three developmental stages.

### UTR and splice junction analysis

For each coding transcript in the *P.falciparum* genome, we calculated the counts for the duplex RNA-seq at each position in the 5’UTR, 3’UTR as well as 100bp downstream of the start codon and 100bp upstream of the stop codon. Since the UTRs vary drastically in size (data not shown), we normalized the UTR lengths into two bins of 50 and 500. All genes with UTRs shorter than 50bp were not considered in this analysis. The threshold of 500 was selected as it is the median UTR length. All UTRs shorter than 500bp were normalized to 50 bins while all UTRs longer than 500bp were normalized to 500bp. We then calculated the normalized expression ratio calculcated against the maximum duplex RNA-seq expression that came from each position in the binned UTR and the 100bp down/upstream coding regions. This was done to study the relative distribution of duplex RNA-seq reads at the 5’ and 3’ gene ends. As low gene expression can make the trends noisy due to their stochastic distributions, we only considered the top 20% genes with UTR expression (~850-1200 genes across the six samples). For each stage, the replicates were averaged for the final analysis. A similar analysis was done for the splice junctions by studying 50bp flanking the top 20% expressed splice junctions.

### Differential expression analysis (GO/KEGG)

Count data was generated for all duplex RNA-seq datasets using the annotated transcripts from (Otto et al. 2010) using featurecounts as described above. Differential gene expression was performed using DESeq2 v1.26.0 in R v3.6.2. The differentially expressed genes across all three stages (adjusted p-value<0.05) were used for GO analysis (198 genes), which was performed using GO terms from PlasmoDB and topGO v 2.38.1 package in R. The ‘Molecular Function” ontology was used.

### *P. falciparum* gene expression (RNA-seq) data

Fastq files of mRNA-seq data was downloaded from the previously published (Otto et al. 2010) dataset. The data was run through the same bioinformatics processing as described about for duplex RNA-seq. The resulting gene counts were normalized using DESeq2 and used for comparison.

### Antisense RNA immunoprecipitation with J2 antibody

For immunoprecipitation of total RNA, 50 μg of RNA from asynchronous, mix stage *P. falciparum* culture was incubated overnight at 4 °C in the reaction buffer containing 15 mM Tris·HCl (pH 7.5),150mM KCl, 5mM MgCl_2_ and 0.5% Triton X-in presence of RNase inhibitor with 10 μg J2 antibody (Scicons). Then, 50 μL of washed (three times in reaction buffer) protein G-Dynabeads magnetic beads were added to the solution of RNA and J2 antibody (ThermoFisher) and then incubated for 4 hours at 4 °C. Complexes were washed four times in reaction buffer and RNA was recovered by phenol-chloroform extraction and ethanol precipitation. RNA integrity, quality and quantity were measured by Qubit 4 fluorometer (ThermoFisher).

### Northern blotting

2ug of J2 immunoprecipitated RNA for each sample was resolved on 6% polyacrylamide gel containing 7 M urea, 90 mM Tris-Borate, and 2.5 mM EDTA (TBE) (pH 8.3) after heating at 95°C for 3 min and instantly chilling on ice. RNA from the gel was subsequently transferred onto N+ nylon membranes (Amersham) using 300 mA constant current for 4 h at 4°C and then cross-linked onto the membrane by UV light. Northern hybridization was done in 0.5 M sodium phosphate buffer (pH 7.6) with gamma-32P-ATP labeled specific oligodeoxyribonucleotides at ~5 × 10^6^ cpm/mL. Following hybridization, membranes were washed twice in 2× SSC, 0.1% SDS at hybridization temperature and once at 0.1× SSC, 0.1% SDS at room temperature for 15 min each and exposed to a PhosphorImager.

### RNA-SHAPE analysis

For SHAPE modification of *P. falciparum* U3 RNA, 5ug of *in vitro* transcribed RNA was denatured by heating at 95^0^C for 2 min followed by immediate incubation in ice for 3 min. Modification was carried out by incubating the RNA in 3X SHAPE buffer consisting of 333mM of NaCl, 333mM of HEPES and 20mM of MgCl 2 at 37^0^C for 30 minutes prior to the addition of 100mM of 2-methylnicotinic acid imidazolide (NAI) for 15 min at 37^0^C. As a negative control, equivalent volume of DMSO was added to the RNA. Based on the modifications obtained from our denaturing gel experiments, we derived secondary structure of *P. falciparum* U3 RNA, based on previously established model (Chakrabarti et al. 2007) using RNAstructure (Xu and Mathews 2016).

## Supplemental Material

Supplemental material is available for this article.

## Acknowledgements

We thank Rania Elbarki, Dipendra Gautam, Arundhati Mohanta and other past and current members of the Chakrabarti laboratory for help with experiments and data analysis. This work was part of DRA’s undergraduate honors thesis project, as part of UNC Charlotte Honors Program in Biological Sciences. This work was supported by UNC Charlotte Faculty Development Award and a consortium award by DSF Charitable foundation to KC.

## Author Contributions

Conceived and designed the experiments: DRA, AO, SK, KC., Performed the experiments: DRA, AO, TB, BZ, AD, SB, and KC. Analyzed the data: DRA, TB, SK, KC. Wrote the paper: SK and KC.

## Supplementary Figures

**Supplementary Figure 1:**
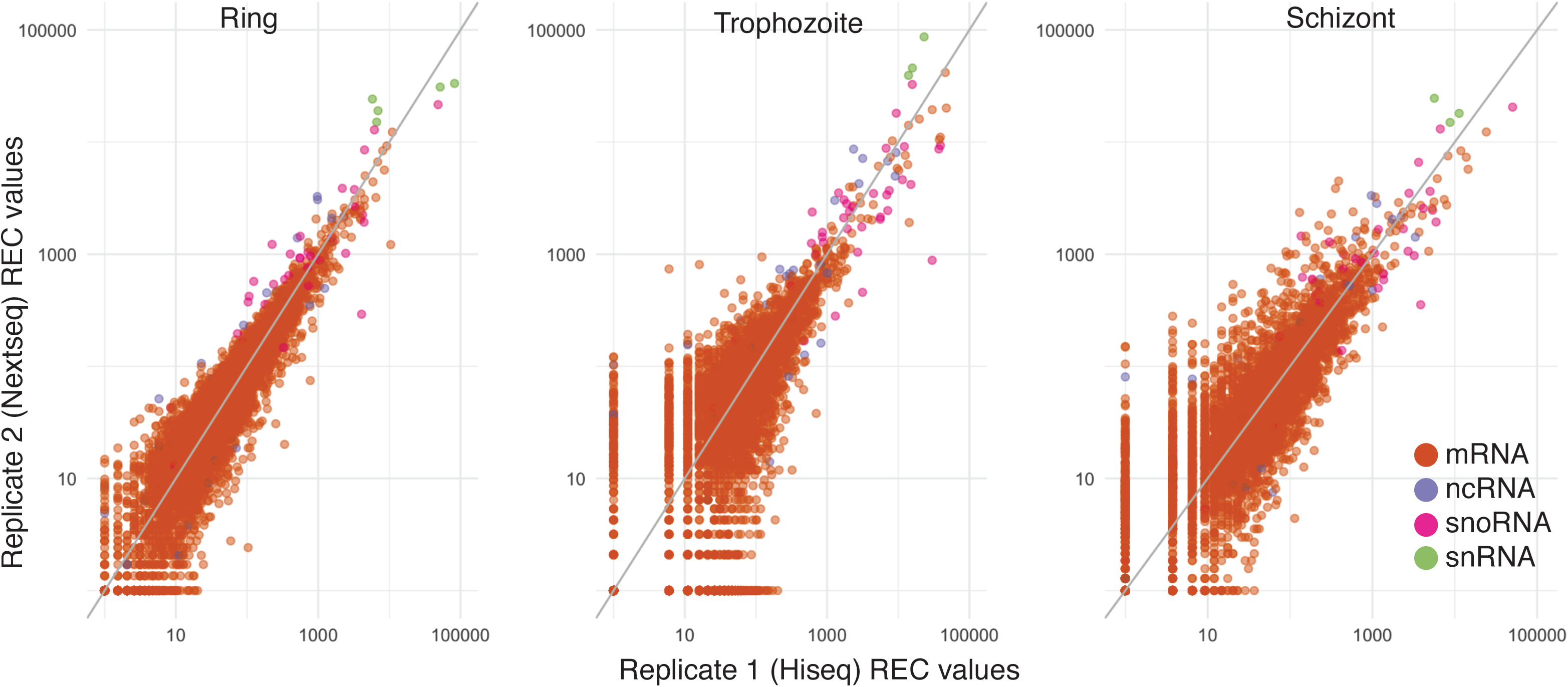
Correlation scatter plots between the biological replicates of the three time point collections of *P. falciparum* after normalization of gene counts by sequencing depth (REC) (see Materials and Methods). The x-axis shows the normalized gene counts for replicates sequenced on HiSeq 2500 while the y-axis shows the normalized gene counts for replicates sequenced on NextSeq500. The axes are plotted in log10 scale and a pseudo count of one is added to all gene counts. The plots show all coding and non-coding genes (5386 genes). Both tRNA and rRNA reads were discarded prior to analysis. The genes are colored by type as indicated: *orange*: mRNA; *blue*: ncRNA; *pink*: snoRNA; *green*: snRNA. Pearson correlation R^2^: 0.87, 0.63, 0.75 respectively.

**Supplementary Figure 2:**
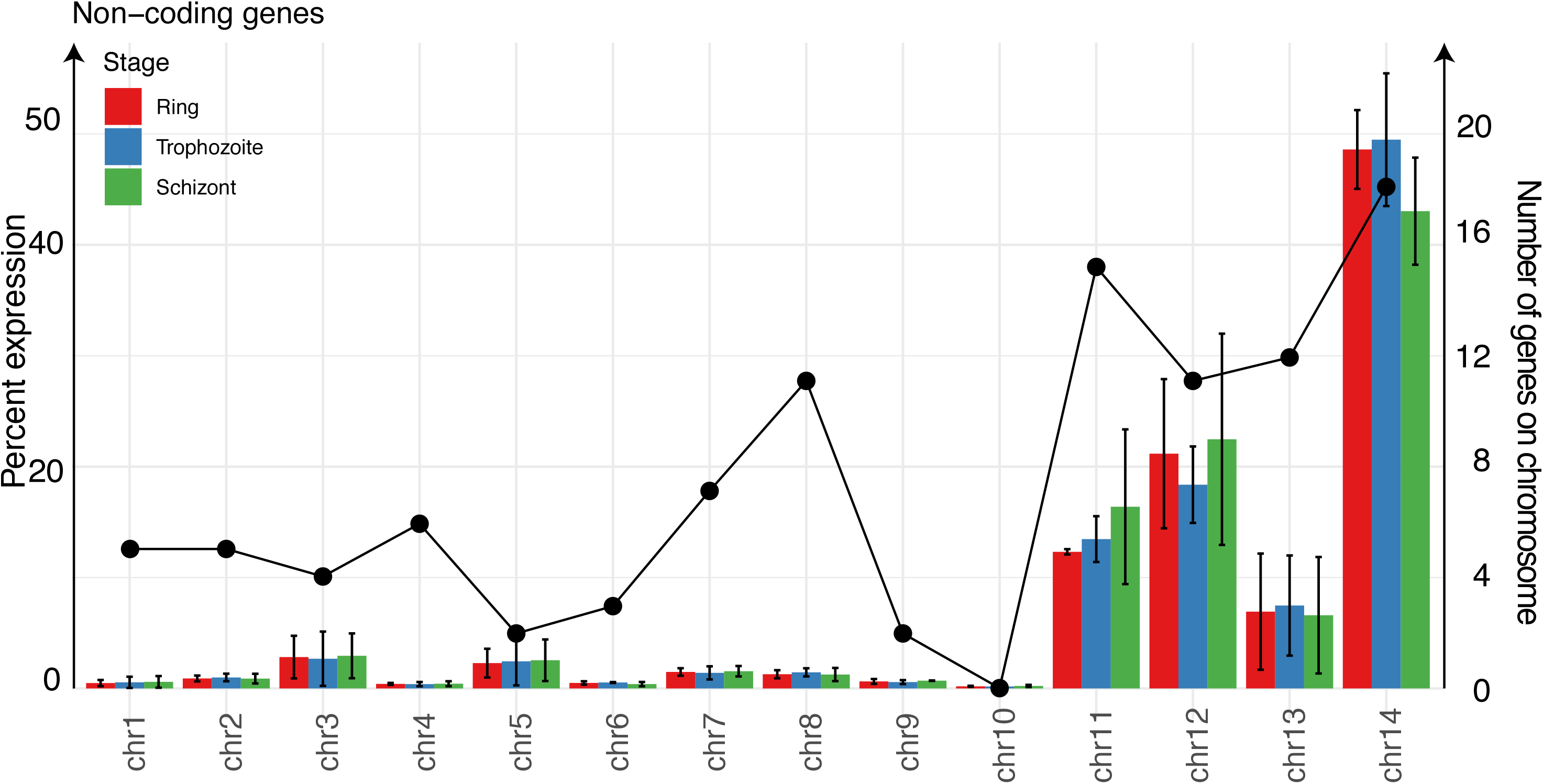
Distribution of uniquely mapped reads mapping to annotated non-coding transcripts in the *P. falciparum* genome. Reads mapping to tRNA and rRNA genes were not included in this analysis. The x-axis shows the 14 chromosomes and the y-axis on the left shows the relative percentage of non-coding reads across each of the chromosomes. The error bars represent the standard deviation across the two replicates of the same stage. The y-axis on the right represents the number of non-coding genes annotated on each chromosome (Chappell et al. 2020). This is shown by the black dots and line on the plot. Colors represent the stage *red*: Ring; *blue*: Trophozoite; *green*: Schizont

**Supplementary Figure 3:**
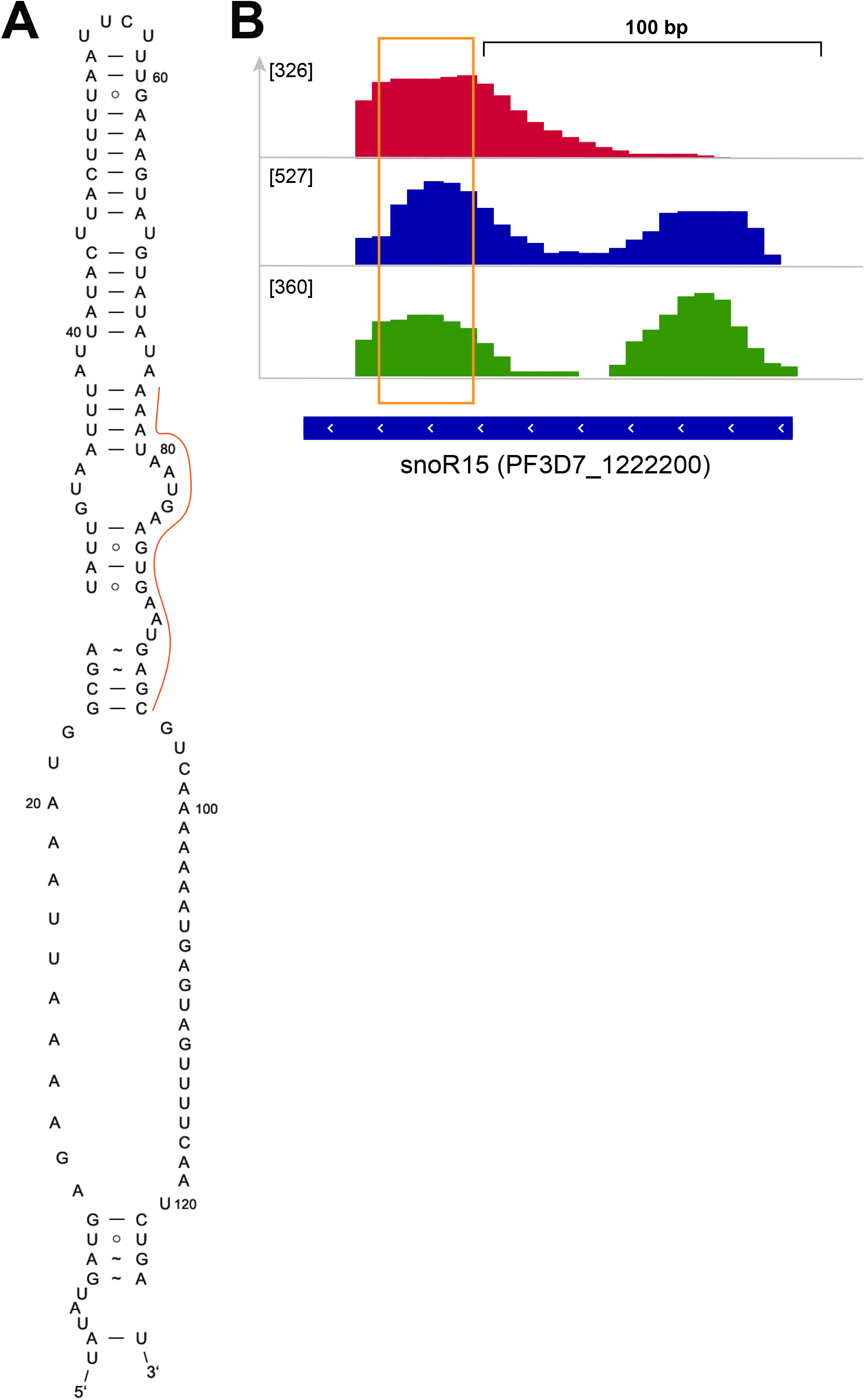
Duplex RNA-seq data for snoR15. (A) Secondary structure of the PfsnoR15 based on in-gel SHAPE modification of mix stage RNA using NAI (B) IGV screenshot of the snoR15 peaks representing duplex RNA-seq read enrichments in three different P. falciparum RBC stages.

**Supplementary Figure 4:**
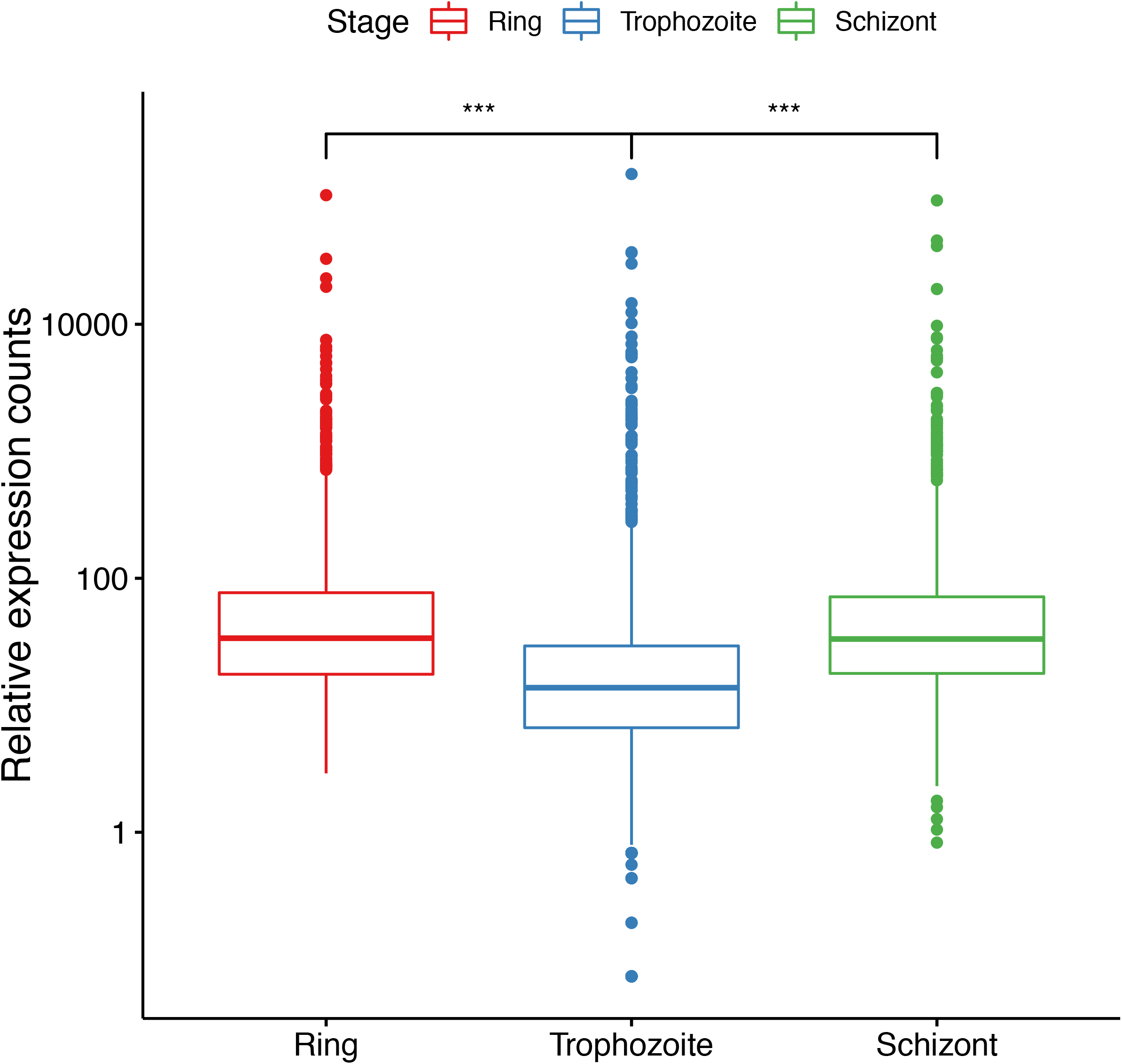
Box plots illustrating the REC values (described in ‘Materials and methods’) of “expressed” coding and non-coding genes (3829 genes) (See Materials and Methods) across the three time point collections of *P. falciparum*. The asterisks indicate a significant difference (two-sided Wilcoxon test, p < 0.001).

**Supplementary Figure 5:**
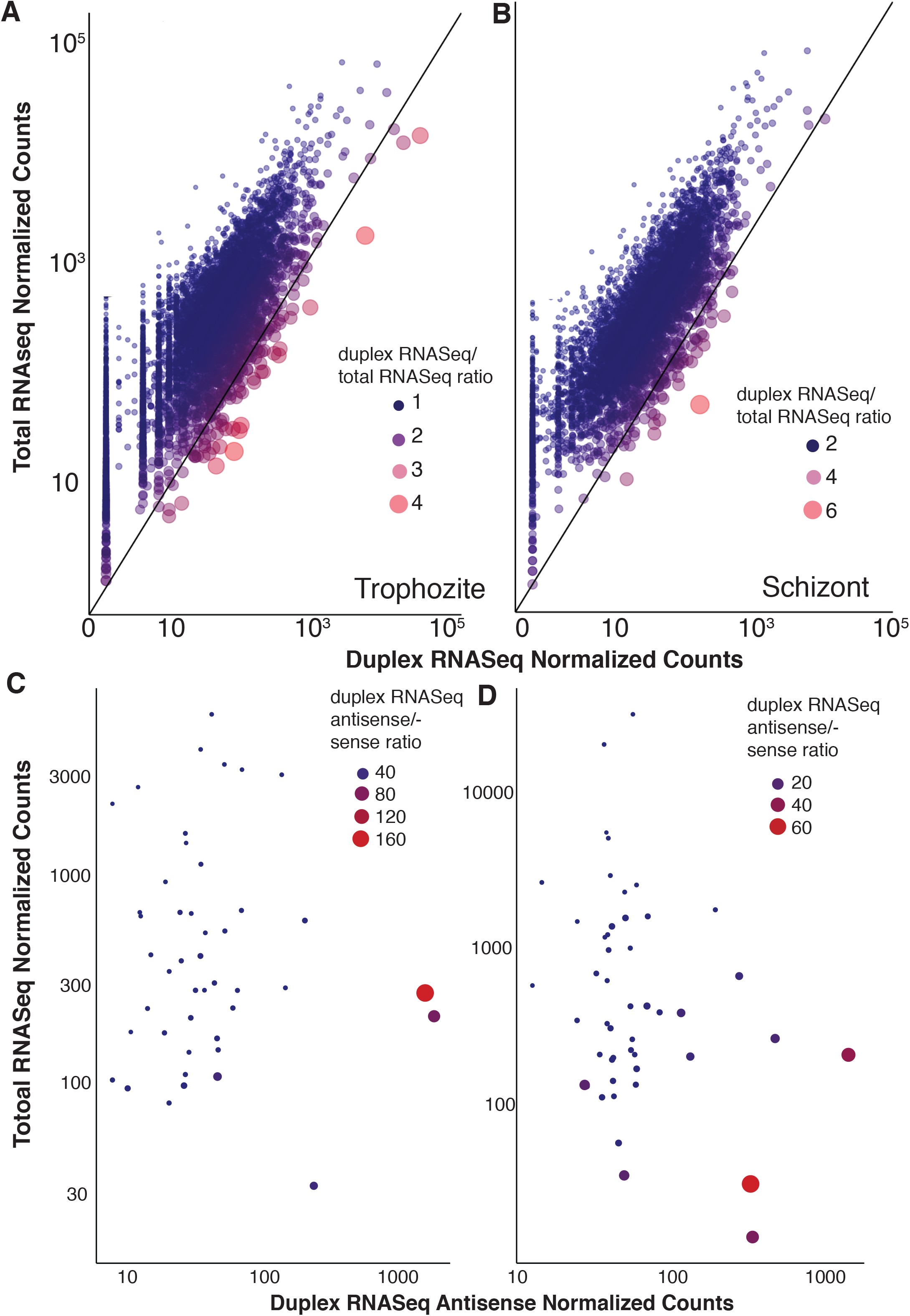
(A-B) Scatter plots for Trophozoite (A) and (B) Schizont showing correlation between normalized gene counts from mRNA-seq and Duplex RNA-seq datasets for Ring stage. A pseudo count of 1.0 is added to the normalized counts for both duplex and mRNA-seq data. The axes are in logarithmic scale. The color and size of the points show the ratio of the duplex RNA-seq normalized counts by the mRNA-seq normalized counts. Larger red points represent genes with higher expression in duplex RNA-seq compared to total mRNA-seq as shown in the legend. (C-D) Antisense duplex RNA-seq counts for the top 50 antisense genes plotted against mRNA-seq counts for (C) Trophozoite and (D) Schizont stages. The size and color of the points indicate the ratio of the duplex RNA-seq antisense to sense counts. Larger red points indicate genes that have more antisense signal than sense signal. Few selected genes with very high and very low ratios are highlighted. A pseudo count of one is added to the normalized counts and axes are in logarithmic scale.

**Supplementary Figure 6:**
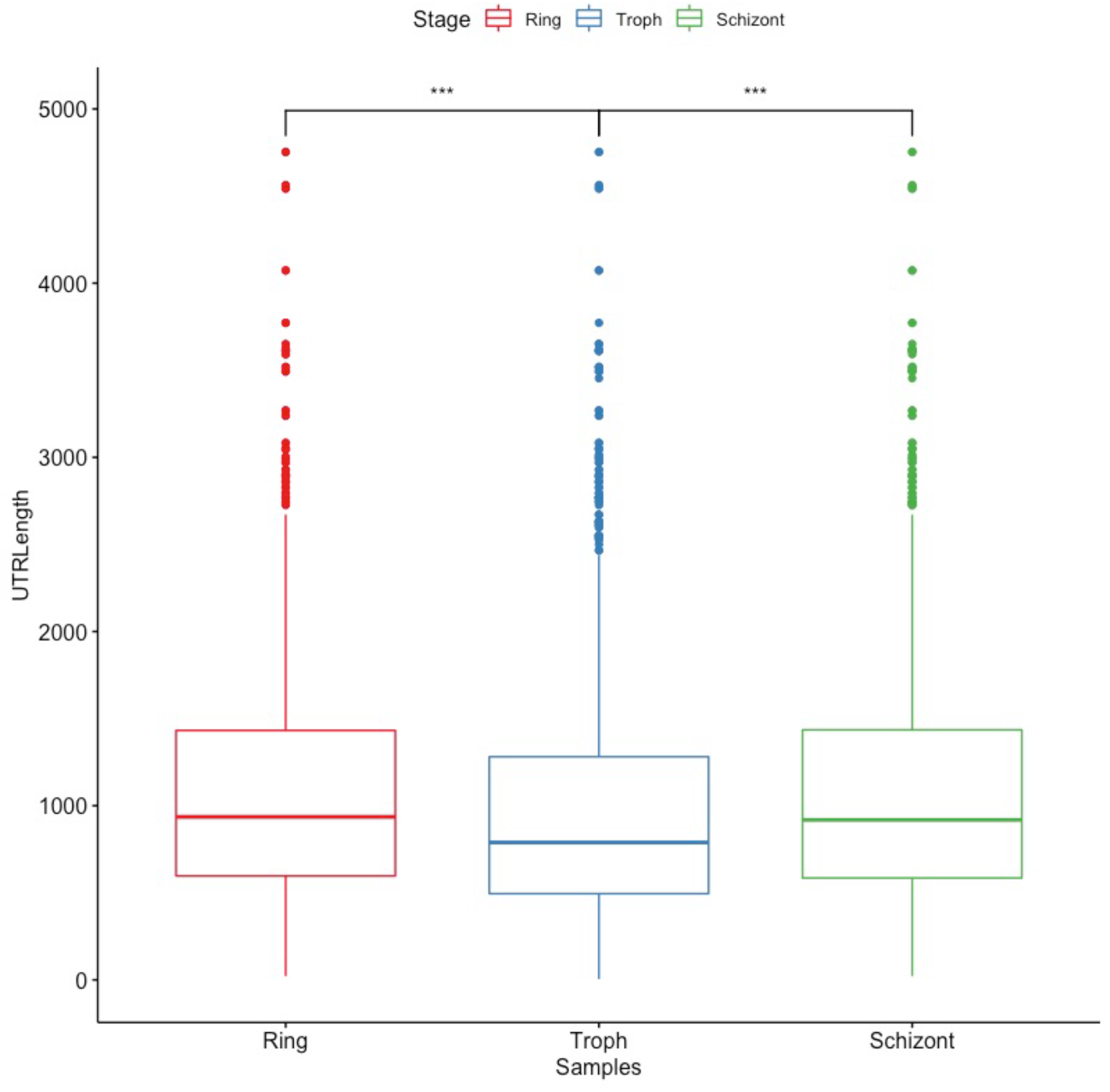
Box plots illustrating the lengths of the 20% most expressed values of “expressed” coding and non-coding genes (3829 genes) (See Materials and Methods) across the three time point collections of *P. falciparum*. The asterisks indicate a significant difference (two-sided Wilcoxon test, p < 0.001).

**Table S1:**
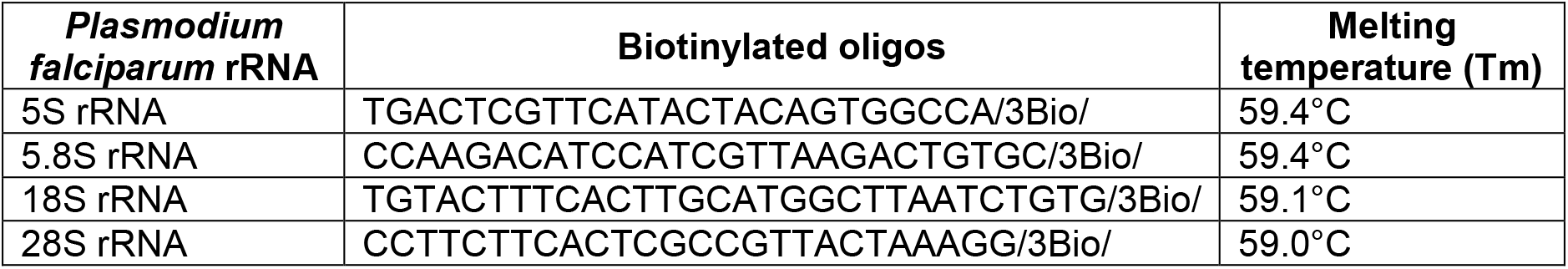
HPLC purified 3’ Biotinylated oligos (3Bio) used for the depletion of *Plasmodium falciparum* ribosomal RNA (rRNA) species.

**Table S2.** Summary of sequence reads obtained from duplex RNA seq libraries. The %total bases Q>=30 and GC content are calculated post-trimming adapter sequences.

| Library | Read length | Total reads | %Total Bases Q<sub>≥</sub>30 | GC Content (%) | Median Insert Size |
| --- | --- | --- | --- | --- | --- |
| *Duplex RNA-seq-1* |  |  |  |  |  |
| Ring | 125X125 | 67,970,256 | 75.89 | 42.25 | 45 |
| Trophozoite | 125X125 | 17,691,860 | 75.03 | 42.82 | 49 |
| Schizont | 105X105 | 10,201,818 | 81.45 | 43.43 | 45 |
| *Duplex RNA-seq-2* |  |  |  |  |  |
| Ring | 75X75 | 178,562,120 | 88.76 | 51.49 | 30 |
| Trophozoite | 75X75 | 437,153,310 | 88.32 | 50.41 | 33 |
| Schizont | 75X75 | 224,498,038 | 89.64 | 51.27 | 30 |

**Table S3.**
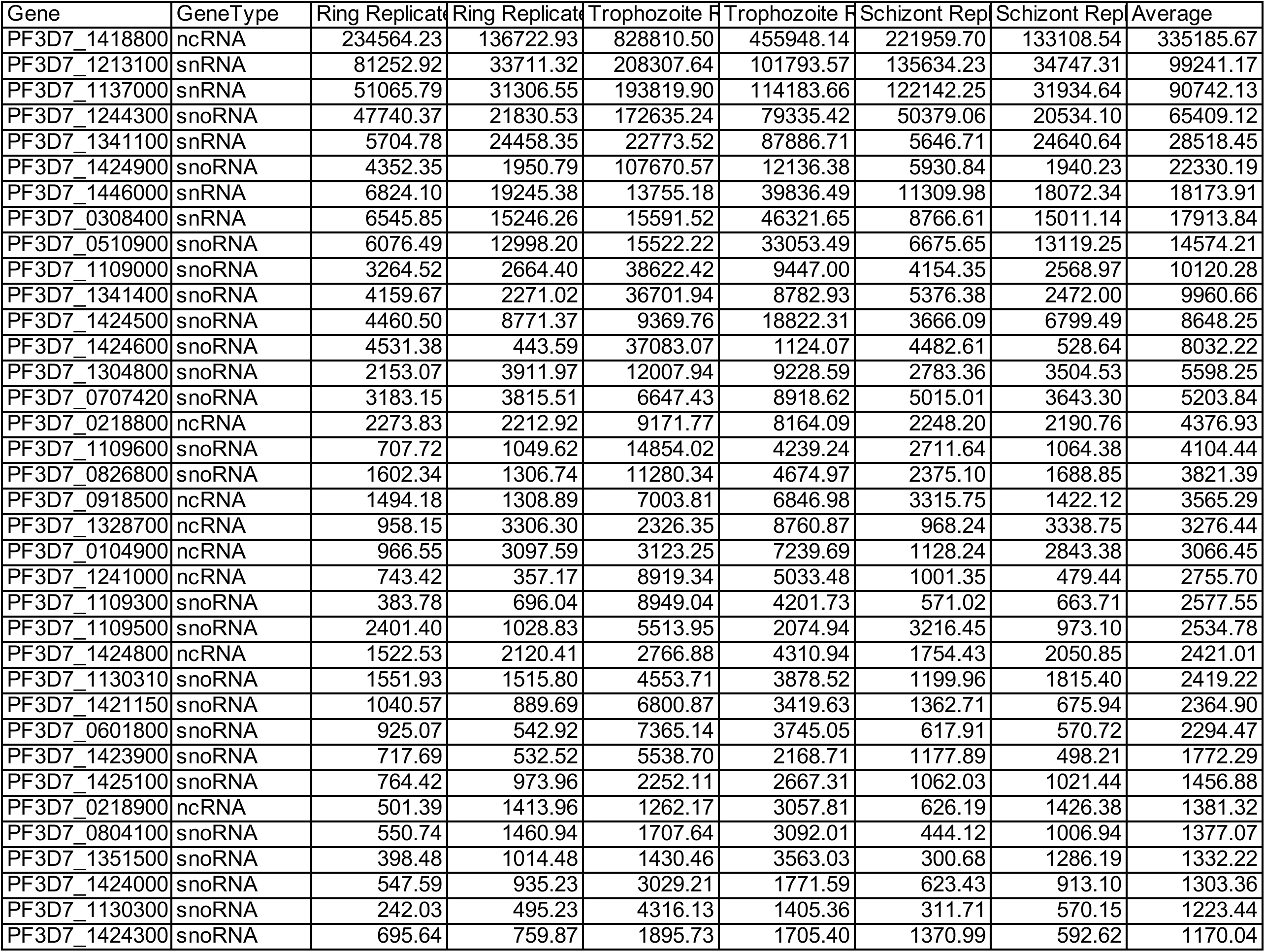
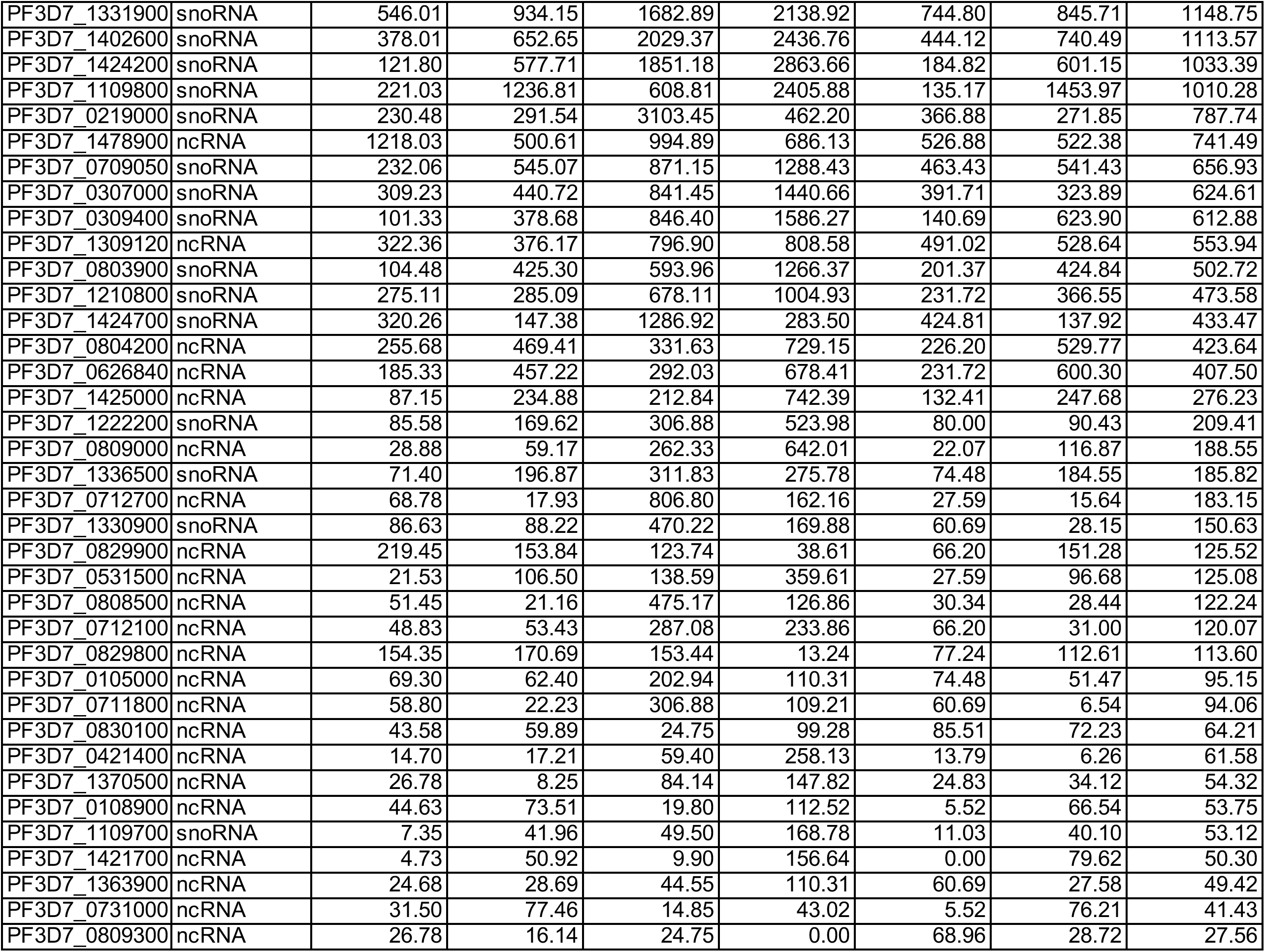
Normalized gene counts of 73 highly base-paired segments of known, conserved structural RNAs

